# Effect of Glycosylation on the Free Energy Landscape of the Catalytic Domain of Human Carbonic Anhydrase IX

**DOI:** 10.64898/2026.08.25.747051

**Authors:** Ritwika Dey, Dulal Mondal, Debayan Chakraborty, Srabani Taraphder

## Abstract

N-linked glycosylation is known to modulate the catalytic function of human carbonic anhydrase (HCA) IX, yet its influence on the underlying free-energy landscape remains largely unexplored. In the present work, we combine extensive all-atom molecular dynamics simulations with kinetic transition network analysis to investigate the effect of glycosylation on the conformational organization of the catalytic domain of HCA IX in both monomeric and dimeric forms. The multidimensional conformational space is discretized into distinct free energy minima using the distribution of reciprocal interatomic distances (DRID), and the effective barriers separating them are estimated using the max flow-min cut formalism. The corresponding free energy landscapes are visualized in terms of disconnectivity graphs, which provide a faithful representation of underlying kinetics. Minimum free energy paths, mean first passage times, as well as frustration metrics are computed to further quantify the effect of glycosylation on landscape topography. Unglycosylated systems are found to exhibit predominantly funnel-like landscapes, with a limited number of metastable states in the vicinity of the native protein fold. In contrast, glycosylation enhances landscape complexity, resulting in a wide array of relaxation timescales. Strikingly, the two glycan chains affect the landscape topography in distinct ways, despite having closely matching sequences. Dimerization couples the glycan chain dynamics, with transitions between key metastable states involving coordinated motions of both the chains. Our work illustrates that interpretation in terms of disconnectivity graphs and transition networks could reveal important insights into the organization of glycoprotein energy landscapes.

## I. INTRODUCTION

A central objective of research in enzyme catalysis is to elucidate the structural and dynamical principles governing their biological functions.^1,2^ The thermodynamic and kinetic behavior of enzymes are known to depend critically on their properties, such as conformational flexibility, binding affinities, folding kinetics, and self-assembly pathways that, in turn, are encoded within the distribution of minima and intervening barriers on the underlying free energy landscapes.^3–9^ In this article, we aim at interpreting the free energy landscape of a glycosylated dimeric enzyme in water in terms of disconnectivity graphs^10–12^ and use it to explore the relevance, if any, of diverse accessible conformations of the enzyme to its catalytic function.

Human carbonic anhydrases (HCAs) are zinc containing metalloenzymes that catalyze the reversible hydration of carbon dioxide to bicarbonate and a proton, a fundamental reaction involved in maintaining physiological pH homeostasis and regulating diverse metabolic processes in the human body.^13–16^ Among the twelve catalytically active human isoforms, HCA II and IX exhibit the highest catalytic efficiencies, with turnover rates of approximately *k*_*cat*_ ~10^6^ *s*^−1^.^17–21^ Owing to its high catalytic activity, monomeric architecture, and ubiquitous expression in normal tissues, HCA II has long served as a prototypical model for elucidating the catalytic mechanism and conformational dynamics of the HCA family. In contrast, HCA IX, the focus of the present study (shown in Figure 1a), is a trans-membrane glycoprotein that catalyzes the same reversible hydration reaction and plays a central role in cellular acid-base regulation.^17,22,23^ Unlike HCA II, HCA IX exhibits limited expression in normal tissues and a marked over-expression under hypoxic conditions in solid malignant tumors.^24,25^ In view of its contribution to extracellular acidification and tumor progression, HCA IX has emerged as a clinically important biomarker and an attractive therapeutic target for cancer diagnosis and treatment.^26–29^

**FIG. 1.**
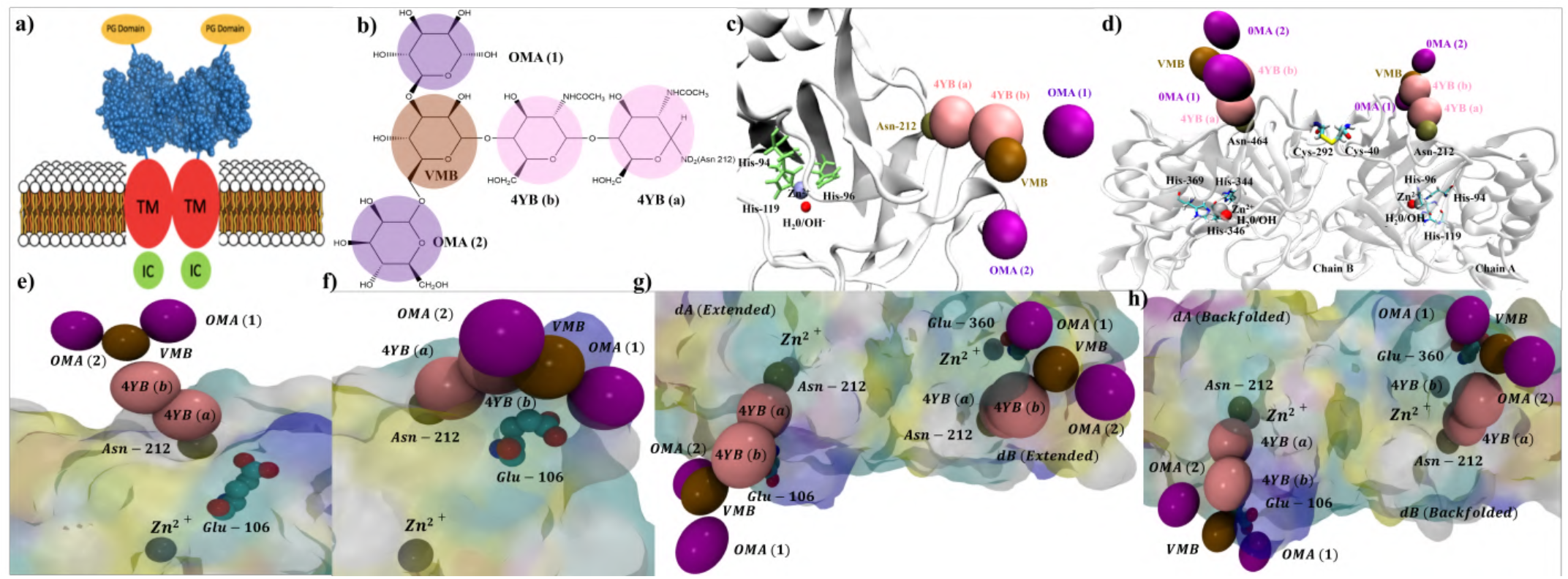
(a) Schematic representation of the trans-membrane, multi-domain structure of dimeric HCA IX, where each monomeric chain (A/B) consists of an N-terminal proteoglycan-like domain (PG, yellow), a catalytic domain (blue), a trans-membrane anchoring motif (TM, red), and a short C-terminal intracellular tail (IC, green). (b) Chemical structure of the glycan chain covalently attached to each chain of the catalytic domain of HCA IX (HCA IX-c). (c,d) The crystal structure (PDB ID: 3IAI^20^) of (c) monomeric (*mA*) and (d) dimeric (*dAB*) HCA IX-c. (e, g) Extended and (f, h) back-folded conformations of the glycan chain attached to (e, f) monomeric (*mA*) and (g, h) dimeric (*dAB*) HCA IX-c. The components of the glycan chain are shown as colored beads: 4YB(a) and 4YB(b) (pink), VMB (brown), OMA(1) and OMA(2) (purple).

In the absence of a fully resolved structure of HCA IX, the present study focuses only on the extracellular catalytic domain for which the crystal structure has been reported (PDB ID: 3IAI).^20^ This domain is highlighted in blue in Figure 1a and designated as HCA IX-c in the remainder of the manuscript. It contains two N-linked glycosylation sites where the N-linked glycan chains are covalently attached to the side-chain amide nitrogen, *N*_*δ* 2_ of Asn-212 (chain A)/Asn-464 (chain B). As illustrated in Figure 1b, the conserved core of each N-linked glycan chain has a branched penta-saccharide structure consisting of (i) two N-acetylglucosamine (GlcNAc) residues, 4YB(a) and 4YB(b), (ii) one *β*-mannose residue, VMB, and (iii) two terminal *α*-mannose residues, OMA(1) and OMA(2). Both glycan chains are found to be oriented away from the protein surface in the crystal structure, as shown in Figure 1c,d. However, owing to the inherent flexibility of the glycosidic linkages and the extensive hydrogen-bonding interactions with the protein surface, both glycan chains are expected to exhibit substantial conformational heterogeneity in solution. For the given structure of the glycan chain, shown in Figure 1b, its conformational space may be broadly spanned by different extended and backfolded structures, presented in Figures 1e-h and classified in Figure 2.

**FIG. 2.**
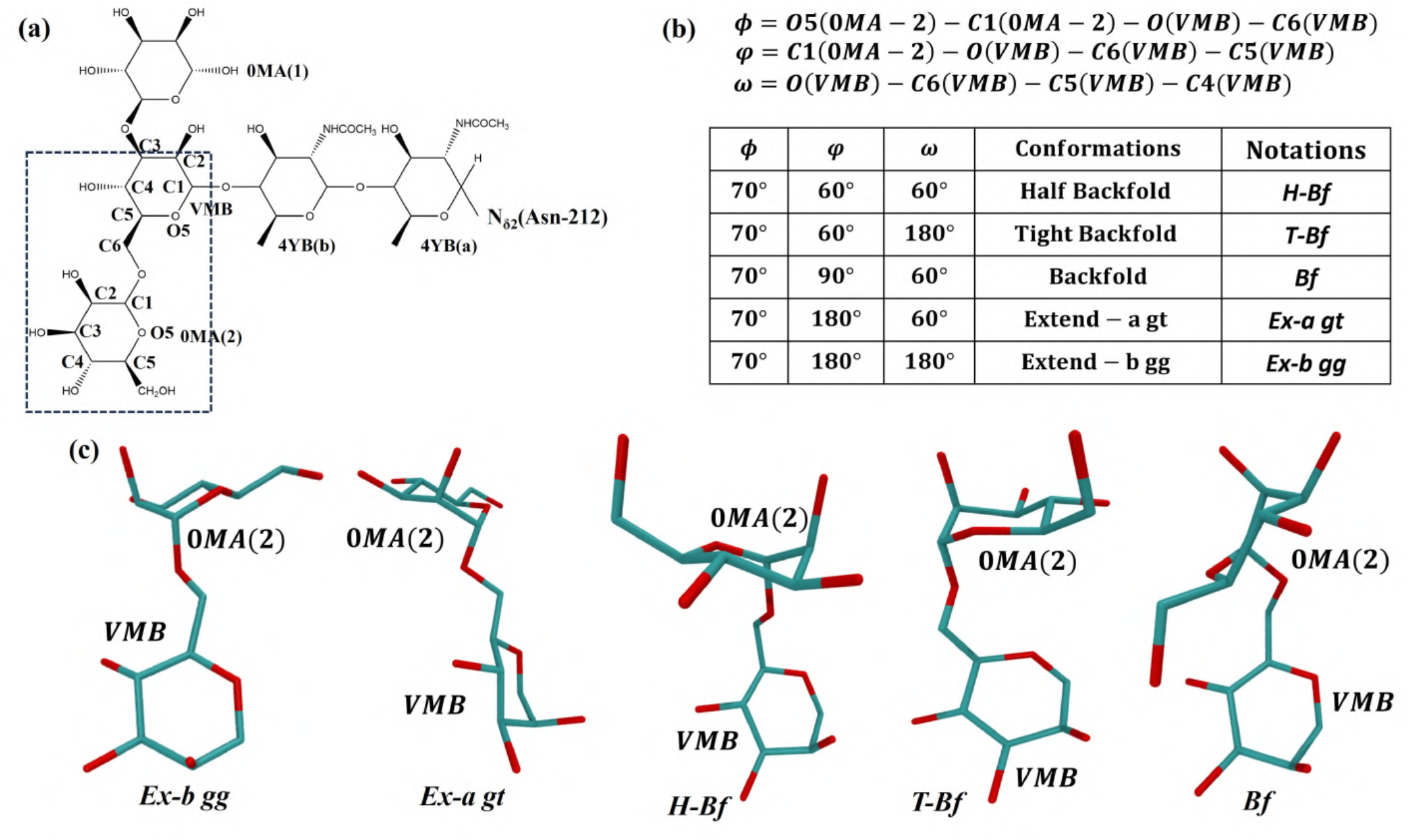
(a) Chemical structure of the glycan chain highlighting the branched VMB and 0MA(2) units, (b) Key descriptors of glycosidic torsional angles, *φ, ψ, ω* and (c) Representative structures of the glycan chain highlighting *Ex-b gg, Ex-a gt, H-Bf, T-Bf* and *Bf* conformations.

Classical molecular dynamics simulations of about 3 *µs* in water, initiated from the crystal structure of glycosylated monomeric HCA IX-c, show that the glycan chain predominantly populates extended conformations that are directed away from the protein surface as shown in Figure 1e.^30^ In contrast, enhanced sampling simulations, based on the accelerated molecular dynamics (aMD) framework, uncovers a substantial population of additional chain conformations, ranging from backfolded to half-backfold and tight-backfold.^30,31^ A representative snapshot of a backfolded glycan chain, derived from aMD simulation of glycosylated monomeric chain A (*mA*) of HCA IX-c in water, is depicted in Figure 1f. As shown in Figures 1e-h, the active site of HCA IX-c includes a catalytically important amino-acid residue, Glu-106 (chain A)/Glu-360 (chain B). A dynamical hydrogen-bonded network (Figure 1f,h) forms between the OMA (1) or OMA(2) terminus of glycan chain and Glu-106 (chain A)/Glu-360 (chain B). Incidentally, the sidechains of these Glu residues participate in the formation of the dynamic hydrogen bonded network through which the rate determining intramolecular proton transfer step is known to take place in HCAs.^17,22,23,31^ The dynamics of water molecules forming this network is found to be correlated to the fluctuations of the distal glycan chains, even when the latter points away from the enzyme surface, in both monomeric and dimeric HCA IX-c. ^31,32^ These observations imply that the conformational plasticity of the glycan chain modulates the activity of the catalytic domain, and may have far-reaching implications in the enzyme’s adaptation to the tumor microenvironment.^24,25^ Interestingly, simulations of dimeric (*dAB*) glycosylated HCA IX-c, also hint at a subtle cross-talk between the two glycan chains, with both adopting either extended (similar to the crystal structure) or backfolded conformations,^32^ as shown in Figures 1g and h, respectively.

As outlined above, our earlier MD simulations primarily focused on detecting thermally accessible conformational variants of the glycan chain in glycosylated HCA IX-c. However, apart from the extended conformation observed in the crystal structure, the existence of other stable conformations of the glycan chain (sampled in enhanced sampling simulations) is yet to be experimentally confirmed. Against this background, a primary objective of the present study is to construct the free energy landscape of monomeric and dimeric HCA IX-c, identify key metastable states in the vicinity of the native-like structures, and determine the transition pathways as well as interconversion rates between key conformations that may be relevant to the catalytic function.

The high dimensionality of biomolecular free energy landscapes often renders their direct visualization impractical. ^12,33–36^ It also complicates the identification of metastable states and their connectivity via transition states. To aid interpretation, numerous linear and non-linear techniques, such as principal component analysis (PCA),^37–40^ time-lagged independent component analysis (TICA),^41,42^ uniform manifold approximation and projection (UMAP),^43^ and t-distributed stochastic neighbor embedding (t-SNE),^44^ which project the high-dimensional landscapes into low-dimensional manifolds, have been proposed. However, these representations can provide a misleading picture of the underlying kinetics unless the ‘perfect’ collective variables (CVs) *aka* ‘reaction coordinates’ are known *a priori*.^45,46^ Although machine learning based approaches have shown great promise in identifying appropriate CVs,^47^ in general, this remains a rather difficult enterprise. To circumvent these difficulties, Becker and Karplus proposed a topographical representation of the landscape, in terms of disconnectivity graphs,^10,33,45^ which preserves the hierarchical organization of the different minima, thereby providing a faithful description of the underlying kinetics. These ideas were developed further by Wales and others,^12,48,49^ and free energy disconnectivity graphs (FEDGs) have since been characterized for a diverse range of systems, from atomic and molecular clusters,^50,51^ to proteins^52^ and nucleic acids,^53^ and glass formers.^54^

In our earlier studies on HCA IX-c in water,^30^ we used a TICA-based method to obtain a low-dimensional representation of the underlying free energy landscape from both conventional MD and aMD simulations. The TICs were constructed from a basis consisting of (i) all pair distances *pd*_*ij*_, between the i^th^ atom of the glycan chain and j^th^ protein atom lying within 5 Å from any atom of the glycan chain and (ii) all 36 (*mA/mB*) / 72 (*dAB*) dihedral angles of the glycan chain(s) as enlisted in Tables S1 and S2. The free energy surfaces projected onto the first two TICs (IC1 and IC2) are shown in Figure S1. These results illustrate how aMD samples the configurational space more extensively than conventional MD, and identifies multiple metastable states corresponding to distinct conformations of the system. Our choice of a large number of physically intuitive collective variables (CVs), as listed above, already reveals the conformational heterogeneity of the glycan chain, and how it is modulated through contacts with the protein surface. However, it is likely that the choice of specific CVs could have influenced our inferences. In the present study, we capitalize on the advantages offered by FEDGs, and provide an unbiased, as well as high-resolution representation of the energy landscapes of HCA IX-c. Furthermore, we quantify how glycosylation affects the landscape topography in the vicinity of the native state, both for the monomeric and dimeric forms of the enzyme.

## II. COMPUTATIONAL METHODS

### A. Classical Molecular Dynamics Simulation

To probe the conformational dynamics of HCA IX-c in explicit solvent under charge-neutralized conditions, we carried out all-atom molecular dynamics simulations using the AMBER18 simulation package.^55^ The AMBER force field was selected due to its extensive validation for glycoprotein systems, particularly through the GLYCAM extension,^56^ which provides a consistent and transferable parameter set for both protein and carbohydrate components. This combination enables reliable modeling of glycosidic linkages, pyranose ring conformations, hydrogen-bonding interactions, and non-bonded contacts in complex glycoconjugates.^57^ A detailed benchmarking of the choice of force fields was performed in our earlier study.^31^ Under identical simulation conditions (including trajectory length), the *ff* 19SB force-field resulted in a substantially broader conformational ensemble for the glycan chain, as opposed to CHARMM36m, which predominantly stabilized extended structures oriented away from the protein surface.^31^ Because back-folded glycan chain conformations promote enhanced exploration of the protein surface and facilitate transient interactions with residues proximal to the active site interactions that may be functionally relevant, the *ff* 19SB^55^ force-field was deemed more appropriate for the present investigation. Accordingly, all monomeric and dimeric simulations were conducted using *ff* 19SB^55^ for the protein in combination with GLYCAM 06j-1^56,58,59^ for carbohydrates. The initial atomic coordinates were obtained from the high-resolution crystal structure of HCA IX-c (PDB ID: 3IAI^20^). The N-linked glycan chain was constructed using the GLYCAM-Web interface,^56,58,59^ and complete system parametrization was performed using the *tleap* module of AMBER18.^55^ The Zn(II)-containing active site was parameterized using the Zinc AMBER Force Field (ZAFF)^60^ to ensure an accurate description of metal coordination geometry and local electrostatics. The water molecules in the solvent box were modeled using the TIP3P force-field.^61^ The details of parametrization are summarized in Table I.

**TABLE I.** Models constructed with *ff* 19SB and GLYCAM 06j-1 force fields for classical MD simulations of glycosylated and un-glycosylated HCA IX-c in water, with the N-glycan linked to Asn-212(chain A)/Asn-464(chain B) in the glycosylated models.

| Model | Description | Force field used |  | Number of water molecules |
| --- | --- | --- | --- | --- |
|  |  | Protein | Glycan |  |
| *U-mA* | Monomeric chain A (mA) without glycan chain | *ff19SB* | – | 17,040 |
| *G-mA* | Monomeric chain A (mA) with glycan chain | *ff19SB* | *GLYCAM06j-1* | 17,120 |
| *U-mB* | Monomeric chain B (mB) without glycan chain | *ff19SB* | – | 16,898 |
| *G-mB* | Monomeric chain B (mB) with glycan chain | *ff19SB* | *GLYCAM06j-1* | 17,100 |
| *U-dAB* | Dimeric (dAB) without glycan chain | *ff19SB* | – | 38,856 |
| *G-dAB* | Dimeric (dAB) with glycan chain | *ff19SB* | *GLYCAM06j-1* | 39,182 |

Each system listed in Table I has been subjected to a three-stage energy minimization protocol consisting of initial steepest descent steps followed by conjugate gradient refinement. The minimized system was subsequently heated gradually to 300 K under controlled conditions. To ensure thorough equilibration of both the protein scaffold and the flexible glycan chain, an extended multistage equilibration protocol was employed.^30,31^ Each system was initially equilibrated in the NPT ensemble for 10 ns, with strong positional restraints of 240 kcal mol^−1^ Å^−2^ applied to residues exhibiting the highest degree of correlated motion within the glycan chain. This stage was followed by 6 successive 10 ns NPT equilibration phases, during which the restraint force constant has been progressively reduced by a factor of two at each stage, reaching a final value of 3.75 kcal mol^−1^ Å^−2^. After complete removal of the restraints, the system was further equilibrated for 200 ns in the NPT ensemble and an additional 200 ns in the NVT ensemble to ensure sufficient convergence of structural and thermodynamic observables. Finally the production runs have been carried out in the NVT ensemble at 300 K, generating 3 *µs* long trajectories for each system.

### B. Identifying free energy minima from MD trajectories using conformational clustering

To partition the conformations sampled along the MD trajectories into distinct free energy minima, we first considered three sets of order parameters (OPs) as described below:

1. Distribution of reciprocal of interatomic distances (DRID): DRID^62,63^ between all the heavy atoms of the protein and the glycan chain.
2. Combination of dihedral angles and pair distances: This set of structural features involves (i) all the 36 (monomer) /72 (dimer) dihedral angles of the glycan chain and the pair distances (*pd*_*ij*_) between any *i*^th^ heavy atom of the glycan chain and *j*^th^ protein residue lying within 5 Å of the glycan chain.
3. Pair distances: *r*_*ij*_ between the *i*^th^ heavy atom of the glycan chain and *j*^th^ protein residue lying within 5 Å of the glycan chain.

To assess the quality of each set, we computed the Variational Approaches in the Markovian (VAMP) scores^41^ using the pyemma code.^64^ The higher VAMP scores for DRID indicate that it retains more of the slow dynamical variance relevant to the Markovian description of the system. Hence, we have primarily employed the DRID metric to identify the distinct microstates (free energy minima) on the energy landscapes.

Since the DRID metric is constructed from reciprocal interatomic distances, it provides a continuous, geometry-based representation that is less sensitive to local angular noise. It also provides an orientation-independent representation of molecular structure and enables efficient quantification of conformational differences without structural alignment.^62^ In this approach, each molecular configuration is encoded by a vector composed of statistical moments of the distribution of reciprocals of interatomic distances. For a set of *n* centroids, the DRID descriptor contains 3*n* components corresponding to the first three moments of the reciprocal distance distribution associated with each centroid. For a given centroid *i*, these descriptors are defined as^62^

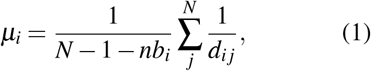

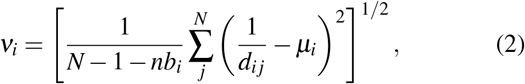

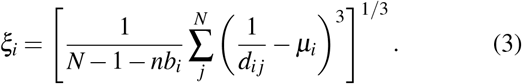

Here, *d*_*ij*_ denotes the distance between *i*^th^ centroid *i* and the *j*^th^ atom; *N* denotes the number of atoms and *nb*_*i*_ is the number of atoms covalently bonded to the centroid *i*. The DRID vector is thus given by^62^

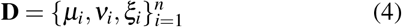

which is invariant under global translations and rotations.^62^ The heavy atoms belonging to non-symmetric groups are selected as centroids to reduce redundancy arising from symmetric atomic environments. The atoms directly bonded to the centroid have been excluded to eliminate the contributions from bond-length fluctuations that do not reflect any meaningful conformational changes. The following four different sets of distances, {*d*_*ij*_} have been employed to construct the DRID descriptors for each of the simulated models.

For the **un-glycosylated systems *U-mA, U-mB, U-dAB***, the sidechain *N*_*δ* 2_-atom of Asn-212 (*mA, mB*) and Asn-212 (chain A)/Asn-464 (chain B) (*dAB*) is connected to a H-atom, instead of the glycan chain. All pair distances between the heavy atoms of the protein (excluding Asn-212 and/or Asn464) and *N*_*δ* 2_-atom of the sidechain of Asn-212 and/or Asn-464 residue are calculated. This model is expected to characterize the intrinsic structural environment of the protein in the absence of any covalently linked glycan chain. The resulting latent spaces have dimensions equal to 4, 652 (*U-mA*), 4, 592 (*U-mB*) and 10, 768 (*U-dAB*).

For the **glycosylated systems *G-mA, G-mB, G-dAB***, the DRID descriptors have been constructed using centroids enlisted in three different ways.

- Focusing *only on the heavy atoms of the protein* (GP), the pair distances have been computed between protein heavy atoms and *N*_*δ* 2_-atom of the sidechain of Asn-212 (chain A)/Asn-464 (chain B), through which the glycan chain is covalently attached within each monomer. This representation is expected to allow a direct quantification of glycan-induced modulation of the local protein environment. The dimensions of the associated latent space, as in the case of un-glycosylated system, are 4, 652 (*GP-mA*), 4, 592 (*GP-mB*) and 10, 768 (*GPdAB*).
- To probe the *glycan-only* part of the glycosylated systems, the pair distances between the heavy atoms of the glycan chain are calculated leading to a latent space spanned by 183 OPs for the models *G-mA, G-mB* and 366 OPs for *G-dAB*. For the dimer, both glycan chains are included in the construction of the DRID descriptors.
- To study the coupling between *both protein and glycan chain* in the glycosylated systems *G-mA, G-mB, G-dAB*, the pair distances have been computed including the heavy atoms of both the protein and the glycan chain(s). The resulting dimensions of the latent space spanned by the respective OPs are 6, 165 (*G-mA*), 6, 063 (*G-mB*) and 12, 225 (*G-dAB*).

The conformational similarity between different configurations defined by the DRID vectors *D* = *µ*_*i*_, *ν*_*i*_, *ξ*_*i*_ and 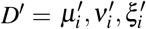 is quantified in terms of the Euclidean distance, Δ(*D, D*^*′*^) given by:^63^

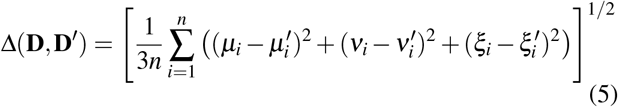

The DRID space was discretized using the regular space clustering method as implemented within pyemma.^64^ The optimal number of clusters identified for each system are tabulated in Table II. The variation of the number of clusters with the distance cutoff is shown in Figure S3. Each cluster, *i*, is considered as a distinct free energy minimum on the energy landscape. The free energy of minimum *i* is expressed as *F*_*i*_ = − *k*_*B*_*T* ln *Z*_*i*_, where *Z*_*i*_ is the partition function. Here, *Z*_*i*_ = *N*_*i*_, where *N*_*i*_ is the number of conformations assigned to cluster *i*. The full partition function, *Z* can be written be written as a superposition sum of the contributions from each local minimum, and is equal to the total number of conformations *N* sampled along the MD trajectory.^10,36^

**TABLE II.** The number of clusters along with their respective lag times, *τ* (ps) for the calculation of the barrier height.

| Models | $\tau$ (ps) | No. of Clusters |
| --- | --- | --- |
| *G-mA* | 50 | 150 |
| *GP-mA* | 25 | 5 |
| *U-mA* | 25 | 4 |
| *G-mB* | 150 | 216 |
| *GP-mB* | 25 | 4 |
| *U-mB* | 15 | 4 |
| *G-dAB* | 150 | 287 |
| *GP-dAB* | 25 | 7 |
| *U-dAB* | 25 | 6 |

### C. Estimation of effective barrier heights and construction of transition networks

The effective rate constant between minima *i* and *j* can be computed using unimolecular rate theory as follows:^10,36^

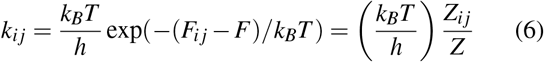

where *F*_*ij*_ is the effective free energy barrier connecting minima *i* and *j*; *Z*_*ij*_ denotes the corresponding partition function; *F* denotes the total free energy, and *Z* the total partition function of the system.

The number of transitions, *N*_*ij*_, between minima *i* and *j* can be directly estimated from our MD trajectories, once the distinct microstates are identified from DRID-based clustering. Following Karplus and Krivov^45^, *N*_*ij*_ is converted into symmetrized edge-capacities, *c*_*ij*_ = (*N*_*ij*_ + *N*_*ji*_)*/*2. Hence, the transitions among various states observed in the MD trajectories can be conveniently mapped into a transition network, where the nodes correspond to the distinct minima connected by edges, whose capacities encode the underlying dynamics of the system. By calculating the maximum flow in this network, the effective barrier between two minima is determined.^10^ The maximum flow between two nodes can be determined from the number of “minimum-cuts” using the Ford-Fulkerson theorem.^65^ To make the computation of minimum-cuts tractable, we used the Gomory-Hu procedure^66^implemented within the *networkx* module.^67,68^

The time-evolution of the probability distribution for the states in the transition network can be described in terms of the linearized master equation:^10^

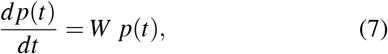

where *p*(*t*) is a vector denoting the probabilities for the system to be in a particular minimum at time *t*, and *W* is the rate matrix.^10^ The transition probability matrix, *T* (*τ*) at lag time *τ* is related to the rate matrix, *W*, by^10^ *T*

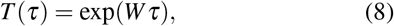

which gives

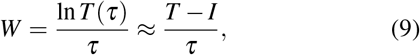

where the matrix element, *w*_*ij*_ = ((*Ñ*_*i j*_*/N*_*i*_) − _*δij*_)*/τ*, with *Ñ*_*i j*_ as the number of minimum-cuts between nodes *i* and *j* obtained using the Gomory-Hu procedure.^66^

The rate constant between minima *i* and *j* is given by

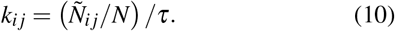

Comparing this expression with Equation 6, and considering that *Z* = *N*, we get *Z*_*ij*_ = *Ñ*_*i j*_ (*h/k*_*B*_*T*) 1*/τ*. Hence, the free energy barrier between minima *i* and *j* is given by: *F*_*ij*_ = −*k*_*B*_*T* ln *Z*_*ij*_. From the databases of minima and barriers, we computed various kinetic observables using the PATHSAMPLE code.^69^ The minimum free energy paths (MFEPs) between selected minima in the transition network were identified using Dijkstra’s shortest-path algorithm.^70^ The mean first passage times (MFPTs) were determined using the new graph transformation procedure.^71^

### D. Visualization of free energy landscapes using disconnectivity graphs

The free energy landscapes were visualized in terms of disconnectivity graphs.^10,33,48^ In contrast to approaches relying on low-dimensional projections, which could lump together minima separated by high barriers, free energy disconnectivity graphs (FEDGs) provide a faithful description of the underlying kinetics. To construct a FEDG, the landscape is partitioned into disjoint basins of attraction by choosing an energy scale that best captures the topological features. Minima within each basin are mutually accessible, whereas inter-basin transitions require surmounting higher barriers. In a FEDG, each vertical line corresponds to a local minimum, originating at its free energy. Two minima are joined at a given level, if the barrier separating them lies below that energy threshold. If the barrier is higher than the threshold, the minima remain disjoint at that level. The FEDG representation, therefore, captures the separation of time-scales inherent in the dynamics of the system. The FEDGs were generated using the disconnectionDPS code.^72^

## III. RESULTS AND DISCUSSION

In the present work, we focus on the construction of the free energy landscape *in the vicinity of the native structure at neutral pH*. Different models of HCA IX-c simulated in this work involve inherently dissimilar properties of the monomeric units of HCA IX-c with distinct chain lengths resulting from different number of constituent amino acid residues: 257 (*mA)*, 252 (*mB*), 509 (*dAB*). To quantify the effect of different conformational states of the glycan chain on the protein conformations and vice versa, while highlighting the difference between the models, each simulated system (*mA, mB, dAB*, as shown in Table I) has been probed systematically to identify (i) conformations of the *protein only* system sampled without (U) and with (GP) glycosylation, (ii) metastable states accessible to the glycan chain only (*gly*) when connected to the protein, (iii) the free energy landscape of the glycosylated HCA IX-c (*G*) in the vicinity of the global minimum and (iv) transition network connecting different metastable states.

### A. Structural Stability of Monomeric and Dimeric HCA IX-c

To investigate the influence of glycosylation and dimerization on the structural integrity of HCA IX-c, the secondary structure elements of the crystal structure were compared with those obtained from the final structures from the 3 *µ*s MD simulations as shown in Figure S4. The difference in the lengths of chains A and B is reflected primarily in the Nterminal region, whereas the catalytic core adopts a highly similar *α/β* fold in both chains. Despite this difference in chain length, the overall secondary structure architecture is well preserved in all simulated systems, indicating that neither glycosylation nor dimerization induces significant alterations to the native fold of the catalytic domain. The central *β*-sheet scaffold remains essentially invariant, while only small-scale fluctuations are observed in a few short helices and *β*-strands. For chain A, as shown in Figure S4(a), the secondary structure elements of the un-glycosylated monomer closely resemble those of the crystal structure. The *α*-helices *α*A, *α*D - H and *α*K are retained throughout the simulations. Likewise, the *β*-sheet network consisting of *β* A - K remains highly conserved. Glycosylation introduces only subtle changes in the boundaries of the short *α*A helix and a few peripheral *β* sheets, whereas the overall topology of the catalytic domain is preserved. A similar behavior is observed in the dimeric systems, where only small adjustments in the lengths of *α*G and *β* G are apparent relative to their un-glycosylated counterpart.

As is evident from Figure S4(b) a comparable trend is observed for chain B. The characteristic *α/β* fold is maintained in both the monomeric and dimeric systems, with the majority of helices and strands exhibiting residue boundaries nearly identical to those of the crystal structure. Minor differences are primarily localized to the *α*A helix and the adjacent *β* A*β* D region, together with small variations in the *β* F, *α*G, and *β* G segments. These changes are confined to the termini of the secondary structure elements and do not propagate to the central *β*-sheet core. The residue-wise root-mean-squared fluctuations (RMSF), shown in Figure S4(c) further demonstrate that glycosylation does not alter the native fold of HCA IX-c, but only induces local structural changes.

### B. Effect of Glycosylation on the Free-Energy Landscape of Monomeric Chains A and B in HCA IX-c

The free energy disconnectivity graphs (FEDGs) for the monomeric chains A and B are depicted in Figures 3 and 4, respectively. A systematic analysis of different contributions to the FEDG of the full system is presented below to reveal how glycosylation progressively reorganizes the conformational landscapes of both monomeric chains while preserving the native fold of the catalytic domain.

**FIG. 3.**
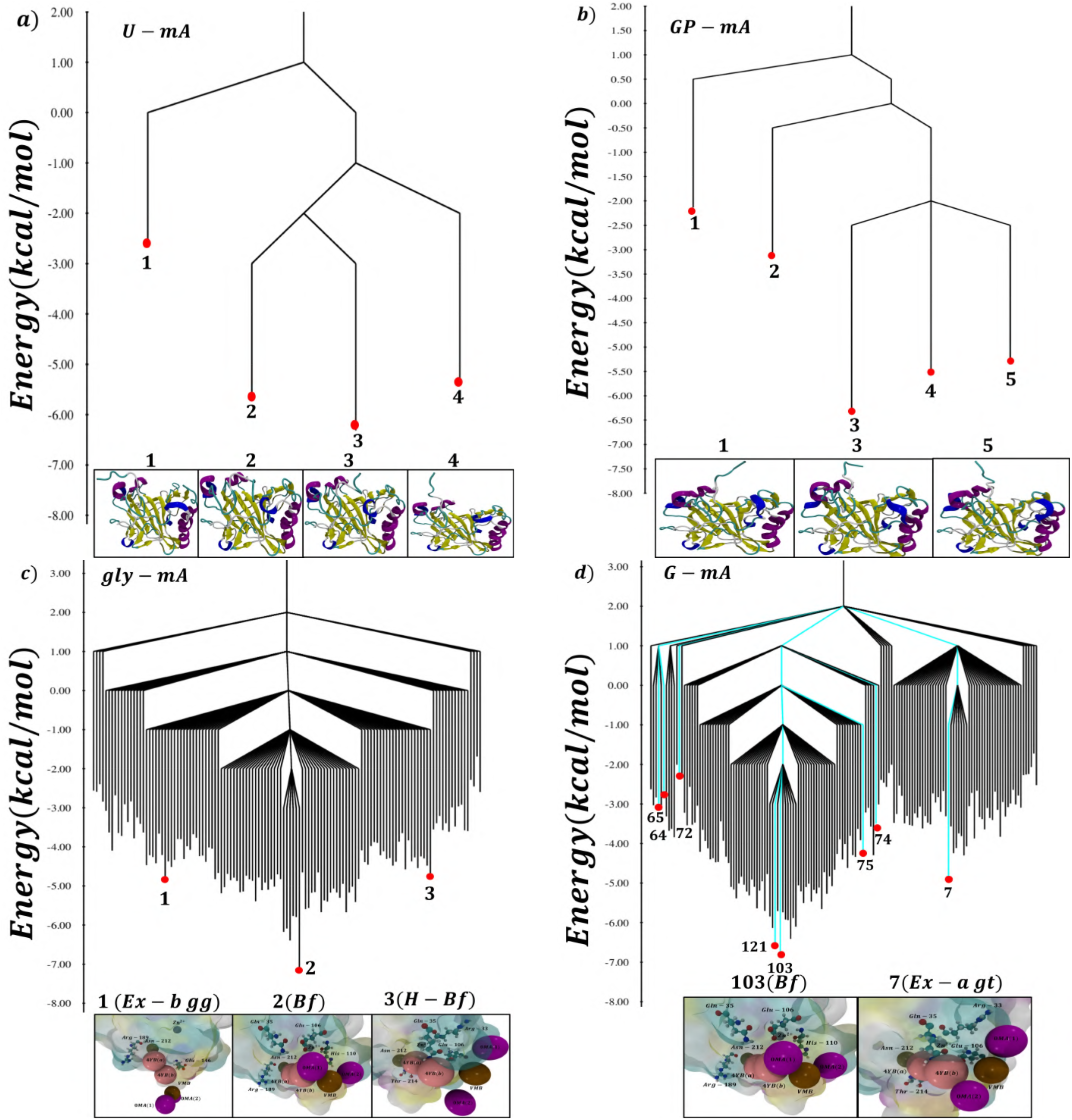
Free-energy disconnectivity graphs (FEDGs) illustrating the free-energy landscapes of (a) *U-mA*, (b) *GP-mA*, (c) the monomeric glycan chain, and (d) *G-mA*. The minima indicated in cyan in panel (d) denote the minimum free-energy pathway (MFEP) connecting the minimum 7 and the global minimum 103 via the intervening metastable states obtained using Dijkstra’s shortest-path algorithm.^70^ The vertical axis corresponds to the free energy in kcal*/*mol.

**FIG. 4.**
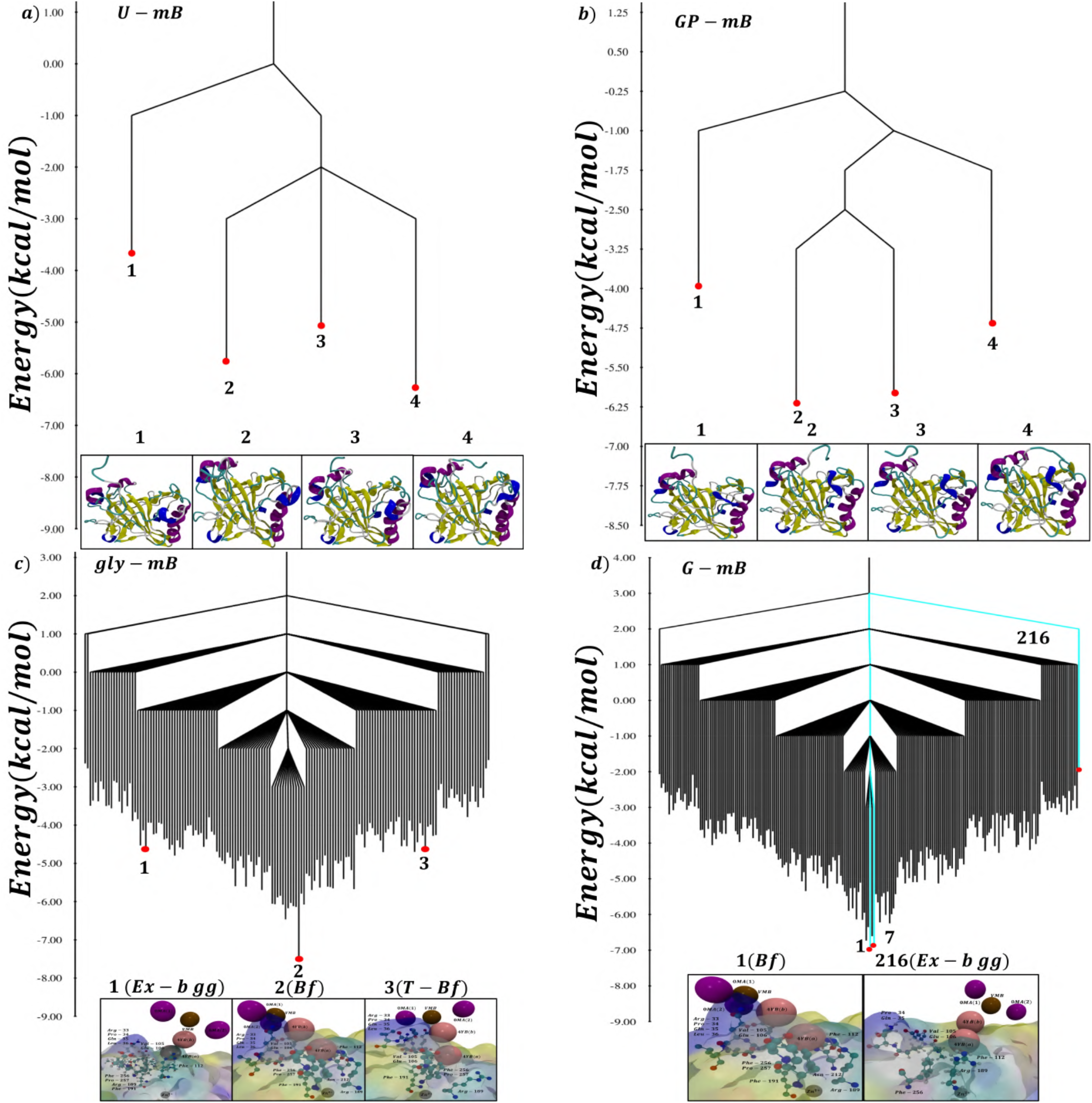
Free-energy disconnectivity graphs (FEDGs) for the models (a) *U-mB*, (b) *GP-mB*, (c) the monomeric glycan chain, and (d) *G-mB*. The minima (cyan) in panel (d) denote the minimum free-energy pathway (MFEP) involved in transitioning of the glycan chain from an extended conformation to a backfolded conformation via metastable states obtained using Dijkstra’s shortest-path algorithm.^70^ The vertical axis corresponds to the free energy in kcal*/*mol.

The un-glycosylated systems, *U-mA* and *U-mB*, exhibit relatively simple funnel-shaped landscapes with only four closely spaced low-energy minima as shown in Figure 3a and Figure 4a. The representative structures associated with these minima preserve the overall structure of HCA IX-c, with structural variations primarily localized to the flexible N-terminal segment and the *α*A helix. The central *β*-sheet core and the remaining secondary structural elements remain essentially unperturbed.

For the *GP-mA* model (in which the DRID basis is constructed using protein atoms only), we find five closely spaced low-energy minima (Figure 3b). The *GP-mB* landscape consists of four low-lying minima (Figure 4b). In both cases, the representative structures maintain the characteristic *β*-sheet architecture of HCA IX-c while displaying enhanced conformational variability in the N-terminal region and the *α*A helix. These observations indicate that covalent attachment of glycan does not significantly expand the conformational ensemble sampled by the protein, or perturb the overall fold.

The FEDGs for *gly-mA* and *gly-mB*, attached to *mA* and *mB*, respectively, are shown in Figure 3c and Figure 4c. For the model *gly-mA*, the DRID basis has been constructed using heavy atoms of the glycan chain only attached to *mA* to obtain (i) their pair distances from each other and (ii) their distances from the *N*_*δ* 2_-atom of Asn-212. The DRID basis for the model *gly-mB* has been constructed similarly using the heavy atoms from their pairwise distances, and their distances from the *N*_*δ* 2_-atom of Asn-464. In both cases, numerous local minima are detected in the FEDGs spanning a broad energy range. This organization of the landscape reflects the intrinsic conformational plasticity of the N-linked glycan, arising from many possible combinations of the glycosidic torsion angles (*φ, ψ*, and *ω*), along with puckering of each sugar ring, which generate diverse extended, tight-backfolded, half-backfolded, and backfolded conformations. Such topologies have previously been characterized for disordered proteins.^73,74^

The FEDGs corresponding to the *G-mA* and *G-mB* models (in which the DRID basis is constructed from both protein and heavy atoms of the glycan chain) are shown Figure 3d and Figure 4d. They subsume the essential features of the glycan only (*gly-mA/mB*) and the protein only (*GP-mA/mB*) landscapes. The representative minima differ not only in localized protein conformations but also exhibit distinct glycan structures. Despite these common features, the overall organization of the *G-mA* and *G-mB* landscapes are significantly different. The FEDG of *G-mA* is characterized by a pronounced double-funnel topology, indicating the presence of low-lying competing minima. This organization is reminiscent of landscapes commonly associated with biological conformational switches,^75–77^ as well atomic clusters exhibiting a low temperature heat capacity peak.^50^ In contrast, the energy landscape of *G-mB* has a single funnel, despite being more ‘rugged’ (i.e. larger number of distinct minima). It is clear that glycosylation substantially increases the conformational heterogeneity of both the monomeric chains of HCA IX-c. However, it reshapes their free energy landscapes in fundamentally different ways, resulting in a double-funneled landscape for *G-mA* and a single-funnel one for *G-mB*.

The origin of the double-funnel topology in *G-mA* may be attributed to the coupling of diverse glycan chain conformations with their neighboring protein residues. The deeper funnel containing the global minimum (node 103) is populated predominantly by the backfolded glycan chain conformation, along with a small fraction of half-backfolded and tight-backfolded conformations. This leads to a rich protein-glycan interaction network whereby multiple backfolded conformations of the glycan chain establish favorable hydrogen-bonding interactions with the protein surface, involving residues Arg-33, Gln-35, Glu-106, Glu-150, Arg-189, Asn-212, Thr-214 and Met-216. In contrast, the competing funnel (leading to local minimum 7) consists of a smaller fraction of backfolded glycan chain conformations. Most of the glycan-protein interactions are localized near the glycosylation site, i.e. Asn-212 (primarily involving Asn-212, Glu-146, Glu-147, Gln-213 and Arg-189). The inter-funnel transition involves a high free-energy barrier of about 7 kcal*/*mol and results from the breaking of persistent intra-glycan hydrogen bonds, the loss of glycan-water hydrogen bonds, and importantly, a reshuffling of the protein-glycan contacts.

### C. Glycosylation in the Dimeric Catalytic Domain of HCA IX-c reshapes the free energy landscape

The FEDGs of the systems *U-dAB, GP-dAB, Gly-dAB* and *G-dAB* are depicted in Figure 5. Together, these FEDGs illustrate yet again the progressive increase in the landscape complexity upon dimerization and glycosylation.

**FIG. 5.**
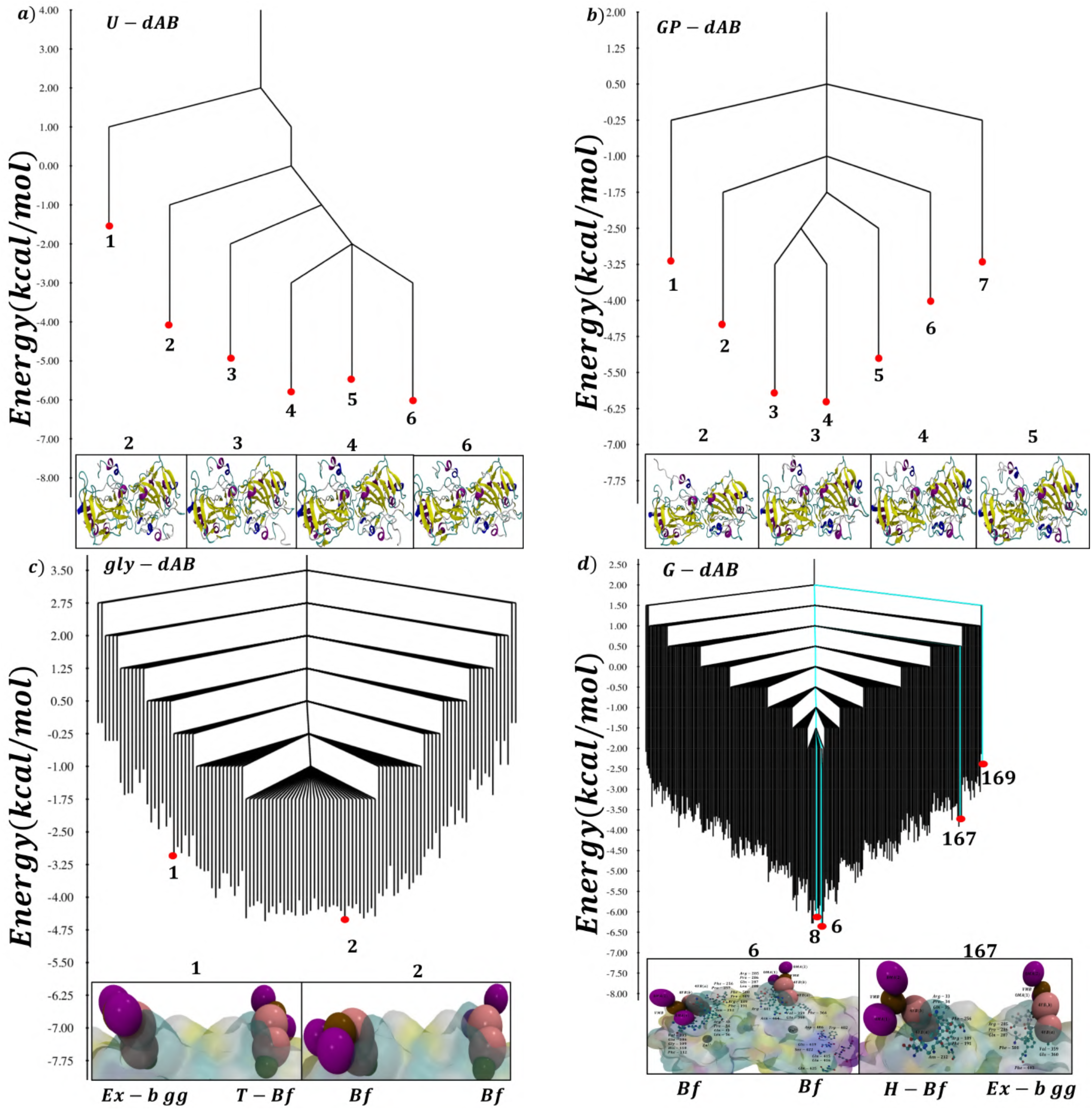
Free-energy disconnectivity graphs (FEDGs) for (a) *U-dAB*, (b) *GP-dAB*, (c) the dimeric glycan chains, and (d) *G-dAB*. In panel (d), the cyan lines indicate the minimum free-energy pathway (MFEP) connecting the extended and backfolded glycan conformations via intermediate metastable states obtained using Dijkstra’s shortest-path algorithm.^70^ The vertical axis corresponds to the free energy in kcal*/*mol.

As is evident from the FEDG (Figure 5a), the landscape of *U-dAB* exhibits a relatively simple funnel, and has six closely spaced low-energy minima. The representative structures preserve the overall dimeric architecture, with the structural differences confined primarily to the flexible N-terminal segments and surface loops, while the catalytic cores of both monomers remain essentially unchanged.

The FEDG of the model *GP-dAB* retains the overall funnel-shaped organization but displays an additional low-lying minimum together with increased branching as shown in Figure 5b. The global fold of the dimer remains largely unperturbed. The representative structures corresponding to the minima 1 − 7 exhibit enhanced structural variability mostly in the flexible N-terminal region while preserving the characteristic secondary structural elements found in HCA IX-c.

The disconnectivity graph corresponding to the glycan chain only system (*gly-dAB*) is shown in Figure 5c. The DRID vectors for *gly-dAB* was constructed from the heavy atoms using their pairwise distances, as well as their distances from both *N*_*δ* 2_-atoms of Asn-212 and Asn-464. This representation isolates the intrinsic conformational preferences of the glycan chains in the dimer, providing a direct view of the accessible sub-states. As is evident, the landscape consists of numerous minima corresponding to extended, half- and tight-backfolded and fully backfolded conformations. Interestingly, the low-lying region is dominated by backfolded states, reflecting yet again that compact glycan chain conformations are thermodynamically favored.

In the *G-dAB* landscape, the dominant funnel characteristic of the protein native state is retained. The large number of local minima primarly results from the glycan chain dynamics. Some representative snapshots corresponding to the backfolded (Bf) and half-backfolded (H-Bf) conformations are shown superimposed on the FEDG in Figure 5d. Overall, our results imply that glycosylation preserves the native dimeric structure, while introducing local frustration. In the global minimum structure, both glycan chains adopt the backfolded conformation (Bf), Half-backfolded (H-bf), tight-backfolded (T-bf) and extended states occupy relatively high-lying regions of the landscape. Some representative snapshots corresponding to the Bf and H-Bf conformations are shown superimposed on the FEDG in Figure 5d. Despite their similar topologies, the dimer landscapes have emergent properties different from those of their monomeric counterparts. From the organization of the local minima, it appears that the glycan chains sample conformations in a correlated fashion, and their dynamics are coupled.

### D. Minimum Free-Energy Pathways (MFEPs)

For the *G-mA* model, the minimum free-energy pathway (MFEP) connecting the global minimum, which is a back-folded state (minimum 103, Figure 3d) to a low-energy extended state (minimum 7, Figure 3d) in the competing funnel was determined using Dijkstra’s shortest path algorithm^70^ (Figures 6a,b). The optimal pathway follows the sequence

**FIG. 6.**
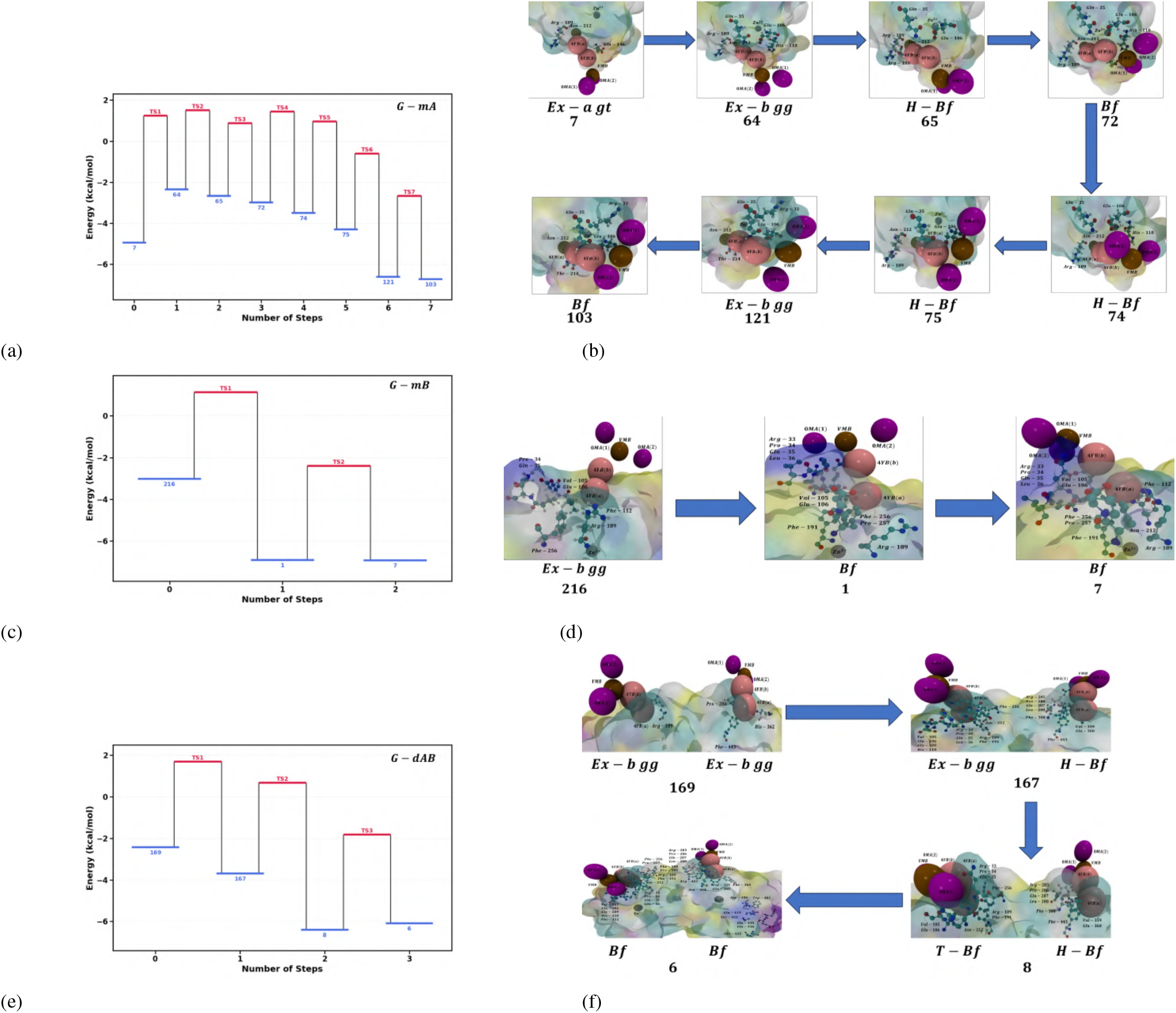
Minimum free-energy pathways (MFEPs) and the corresponding glycan conformational transitions for the glycosylated HCA IX-c models. Panels (a), (c), and (e) depict the MFEPs obtained using Dijkstra’s algorithm implemented in PATHSAMPLE for the *G-mA, G-mB*, and *G-dAB* models, respectively. Panels (b), (d), and (f) illustrate the representative structural intermediates associated with the MFEPs, highlighting the sequential transitions between distinct glycan conformational states.

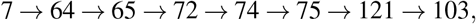

indicating that the transition proceeds through a series of intermediate minima rather than a single barrier-crossing event. Interestingly, the rate-determining step involves a single torsion flip, which results in the conformational transition from the initial *Ex-a gt* to a *Ex-b gg* state. The representative structures associated with these minima illustrate that progression from an extended to the backfolded conformation occurs via several half-backfolded intermediates. The transition rates calculated using the NGT formalism^71^ for model *G-mA* are tabulated in Table III. The substantially faster forward transition indicates that the backfolded conformation is both thermodynamically and kinetically favored.

**TABLE III.** Minimum free-energy pathways connecting an extended glycan state (State 1 or a selected local minimum) to a backfolded glycan state (State 2 or the global minimum) in the glycosylated monomeric and dimeric HCA IX-c models. The corresponding forward (*k*_1 → 2_) and reverse (*k*_2 → 1_) transition rates were calculated using NGT^71^ along the respective MFEPs. The state indices denote the minima defining each pathway.

| Model | State index | | $k_{1 \rightarrow 2}$ ($s^{-1}$) | $k_{2 \rightarrow 1}$ ($s^{-1}$) |
| --- | --- | --- | --- | --- |
|  | State 1 | State 2 |  |  |
| *G-mA* | 7 | 103 | $0.167 \times 10^6$ | $0.67 \times 10^4$ |
| *G-mB* | 216 | 7 | $0.94 \times 10^7$ | $0.23 \times 10^5$ |
| *G-dAB* | 169 | 8 | $0.32 \times 10^7$ | $0.77 \times 10^4$ |

For the *G-mB* model, the MFEP connecting an extended conformation in the *Ex-b gg* state (minimum 216, Figure 4d) to the global minimum, which is a backfolded conformation (minimum 7, Figure 4d) is shown in Figure 6c and 6d. The optimal pathway

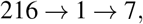

proceeds through a single intermediate minimum. Strikingly, the mechanism does not involve any half backfolded configuration, indicating a more direct transition than that observed for chain A. The transition rates for the *G-mB* model are summarized in Table III. The pronounced asymmetry between the forward and reverse rates demonstrates a strong kinetic preference for the backfolded conformation.

The MFEP for the model *G-dAB* is shown in Figure 6e,f. The optimal pathway connecting a high-energy extended state (minimum 169, Figure 5d) and the global minimum, which is a backfolded conformation (minimum 6, Figure 5d) follows the sequence

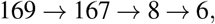

The transition proceeds through two intermediate minima.

The representative structures associated with these minima illustrate the progressive folding of the glycan chains from extended to backfolded conformations as shown in Figure 6f. As is evident from Figure 6f, each elementary step corresponds to a single flip about the *ω* angle for one of the glycan chains, suggesting that multiple flips may involve higher free energy barriers. At no point along the transition pathway, the two glycan chains adopt similar torsional sub-states, further hinting at a non-trivial cross-talk among the chains. The transition rates are summarized in Table III. The asymmetry between the forward and backward rates, demonstrates the persistence of a strong kinetic preference for the backfolded state.

### E. Molecular origin of the highest free energy barriers along MFEPs

As described in the previous section, the highest barrier separating minimum 7 and minimum 64) along the MFEP connecting the in *G-mA* corresponds to an inter-funnel, rather than intra-funnel transition. To gain further insight into the molecular origin of the highest free-energy barriers along the MFEPs, the electrostatic and van der Waals components of the interaction energies between the glycan and protein were examined for local minima immediately preceding and following the rate-determining step. For all three glycosylated systems, the highest barrier is accompanied by a marked enhancement of electrostatic stabilization upon crossing the transition state. In the *G-mA* model, the transition from minimum 7 to 64 results in the electrostatic interaction becoming favorable by about 10 kcal/mol (see Figure S5). However, the van der Waals interaction becomes far less optimal. As a result, minimum 64 is destabilized relative to 7. For the *G-mB* model, the electrostatic interactions become favorable by about 8 kcal/mol during the transition from minimum 216 to 1 (see Figure S5). However, the transition also results in several non-optimal contacts. In the *G-dAB* model, the highest-barrier transition (167 → 8) is associated with a substantially larger electrostatic stabilization (about 22 kcal/mol), which offsets the non-optimal Van der Waals interactions to a greater degree as compared to the monomeric glycosylated system. This enhanced stabilization likely arises from the larger interfacial contact surface and the greater number of favorable glycan-protein interactions available in the dimeric assembly as shown in Figure 6f and Figure S5.

### F. Protein Residues Correlating the Motion of the Glycan Chain

The interaction network between the glycan and protein residues across monomeric (chain A and chain B) and dimeric HCA IX-c systems reveals a hierarchical and state-dependent pattern that governs glycan conformational transitions. Across all systems, distinct glycan conformations, including extended (*Ex-b gg, Ex-a gt*), half-backfolded, tight-backfolded and backfolded states, are stabilized by specific residue clusters distributed along the protein surface. In the extended conformations, the glycan chain predominantly interacts with surface-exposed residues located on the coils and turns, including Arg-33, Pro-34, Gln-35, and Leu-36. Additional contacts with Val-105 and Glu-106 in chain A, and Val-359 and Glu-360 in chain B, indicate that the glycan samples peripheral regions with relatively weak and transient hydrogen-bonding interactions. These interactions are conserved across both monomeric chains, although the spatial arrangement of residues leads to subtle differences in accessibility and orientation. As the glycan chain transitions toward partially folded (half-backfolded) states, the interaction network extends deeper into the protein surface. Residues, such as Glu-106, Phe-112, Asn-212 (*β* I sheet), Arg-189 (*α*G helix), and Phe-191 form a stabilizing hydrogen-bonding network that supports intermediate conformations. In chain B, this transition involves stronger engagement of these residues, consistent with the observed kinetic bias toward backfolded states. In contrast, chain A exhibits a more distributed interaction network, allowing greater accessibility of extended conformations. In the fully backfolded and tight-backfolded conformations, the glycan chain establishes extensive contacts with residues Glu-106, Phe-256, Pro-257, Arg-285, Pro-286, Gln-287, and Leu-288, which collectively stabilize back-folded glycan arrangements. In the dimeric system, additional residues from both chains contribute to this interaction network, including Glu-360, Arg-441, Asn-464, Asp-486, Trp-482, Ser-422, Glu-415, and Gln-425. These interactions indicate that glycan folding is accompanied by deeper insertion and enhanced coupling between the two subunits. A key feature across all systems is the role of Glu-106 (chain A) and Glu-360 (chain B), which act as gatekeepers regulating access of the glycan to the active site residues. Their involvement is particularly pronounced in the dimer, where coordinated interactions across both chains facilitate glycan stabilization. The residue-level analysis demonstrates that glycan conformational transitions are governed by a conserved, yet context-dependent interaction network, progressing from surface contacts in extended states to deeply embedded interactions in backfolded conformations. While the intrinsic interaction pattern is similar across chain A and chain B, differences in residue organization and local environment lead to distinct kinetic behavior. In the dimeric system, this interaction network expands to include residues from both chains, resulting in emergent inter-chain coupling and enhanced stabilization of compact glycan states.

### G. Distribution of Mean First Passage Times (MFPTs)

The mean first-passage times (MFPT) were calculated using the new graph transformation (NGT) algorithm implemented within the PATHSAMPLE code,^78^. The unglycosylated (*U*) models exhibit narrow MFPT ranges, varying from ≈ 800 to 4400 ps. (Table IV), indicating rapid relaxation towards the global minimum. The relatively small spread in the MFPT values suggests that the metastable states are connected through low kinetic barriers, enabling efficient interconversion. Furthermore, the absence of long-lived metastable states is consistent with a predominantly single-funneled topology of the free-energy landscapes. Overall, the *U* models display kinetically homogeneous behavior, characterized by fast and direct relaxation pathways to the global minimum.

**TABLE IV.** Characteristic MFPT timescales spanned by the metastable states to reach the global minimum for the models under investigation. For G-mB, the transition from one of the nodes to the global minimum is extremely fast as compared to the rest, and has been shown separately.

| Models | $\tau_{MFPT}$ (ps) |
| --- | --- |
| *U-mA* | 2035 – 3137 |
| *GP-mA* | 982 – 2500 |
| *G-mA* | 37168 – 6095906 |
| *U-mB* | 824 – 1816 |
| *GP-mB* | 1072 – 2872 |
| *G-mB* | 5898; 99362 – 119540 |
| *U-dAB* | 2530 – 4425 |
| *GP-dAB* | 996 – 1672 |
| *G-dAB* | 5898 – 46595 |

The MFPT ranges for the *GP* models and the fully glycosylated (*G*) models are summarized in Table IV. The *GP* models exhibit relatively narrow MFPT ranges from ≈ 900 − 2900 ps, indicating that the conformational landscape corresponding to the protein subsystem remains kinetically well connected and relaxes rapidly toward the global minimum. In contrast, the fully glycosylated systems display substantially broader MFPT ranges, spanning nearly three order of magnitude. Among them, *G-mA* exhibits the largest kinetic span from 37 − 6095 ns, reflecting the presence of long-lived metastable states, and consistent with the double-funneled topology of the landscape. Although the models *G-mB* and *G-dAB* also show significantly slower relaxation times (5.89 − 119.54 *ns* and 5.89 − 46.59 *ns*, respectively) than their corresponding GP models, their comparatively narrower MFPT ranges suggest less severe frustration. These observations demonstrate that the explicit inclusion of glycan chains markedly increases the kinetic heterogeneity of the free-energy landscape by stabilizing metastable states and slowing transitions toward the global minimum.

The probability distributions of ln(*τ*_MFPT_) provide direct insight into the kinetic organization of the underlying free-energy landscapes. For the *U-mA* model, the narrow peak indicates that relaxation toward the global minimum is governed by a single dominant timescale as shown in Figure 7(a). For the *GP-mA* model, the broader and somewhat asymmetric unimodal distribution indicates that, although relaxation is still dominated by a single characteristic timescale, the landscape exhibits greater kinetic heterogeneity than *U-mA* (Figure 7a). In other words, the glycosylated system samples a wider spectrum of relaxation pathways for reaching the global minimum. These finer features, are however, not immediately apparent from the FEDG shown in Figure 3. The bimodal distribution observed for the *G-mA* model (Figure 7a reveals the existence of two well-separated relaxation timescales, consistent with the kinetic signatures expected for a multi-funnel free-energy landscape.^79^ The first peak corresponds to relaxation within the dominant funnel, whereas the second peak corresponds to the inter-funnel transition. This interpretation is further supported by the distribution of the number of elementary rearrangements, where most local minima require two or eight elementary steps to reach the global minimum (Figure 7c).

**FIG. 7.**
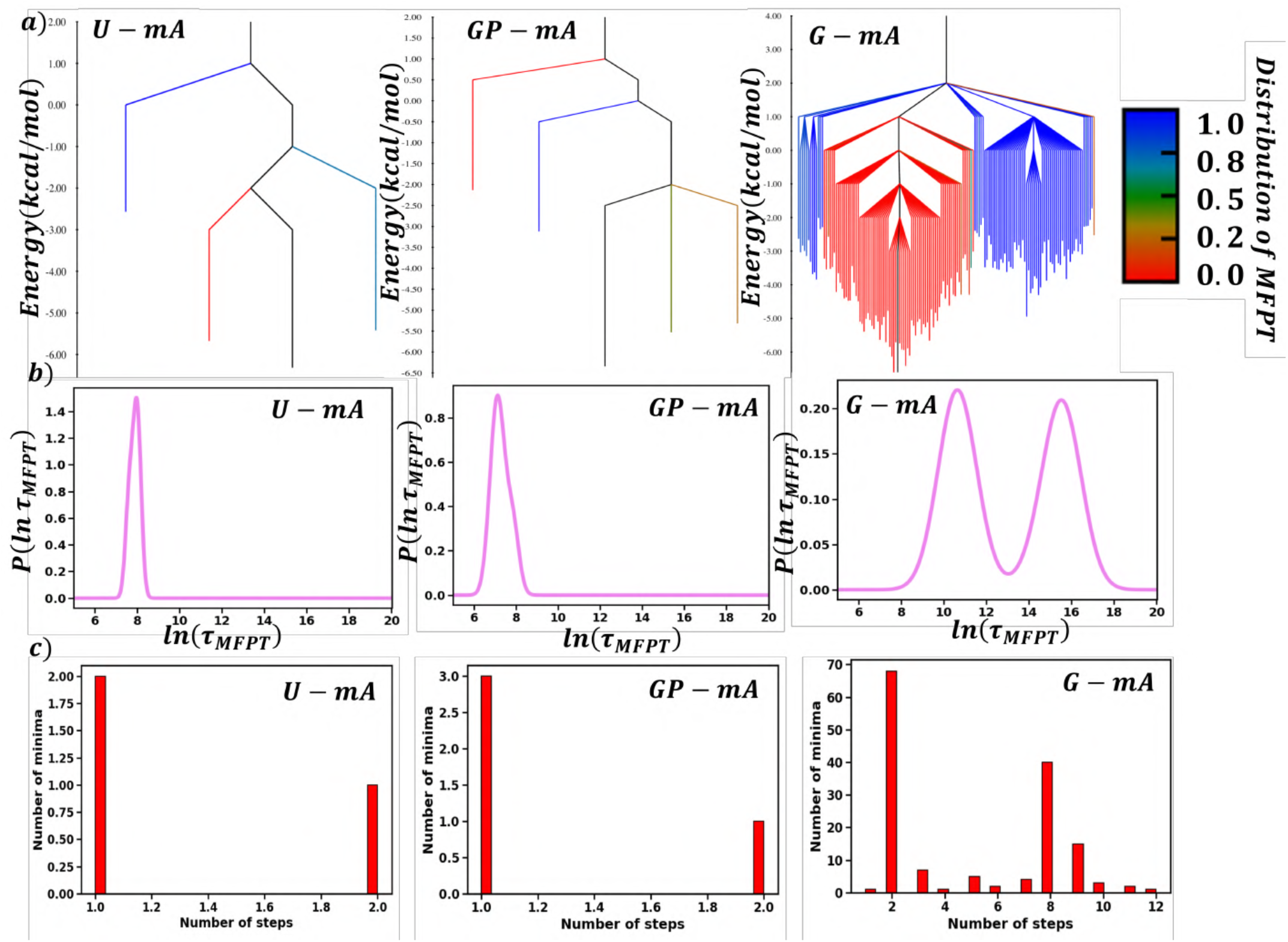
(a) Disconnectivity graphs and (b) probability distributions of the logarithm of the mean first-passage time, *P*(ln *τ*_MFPT_), for *U-mA, GP-mA*, and *G-mA* models. Panel (a) presents the disconnectivity graphs constructed from the local minima. The branches are colored according to the normalized ln *τ*_MFPT_ distribution associated with the corresponding minima, as indicated by the color bar. Panel (b) shows the probability distributions of ln(*τ*_MFPT_) calculated from the MFPT values of all local minima with respect to the global minimum. The distributions characterize the kinetic timescales governing relaxation over the respective energy landscapes, where we obtain bimodal distribution for *G-mA*, validating the double funnel feature in the landscape at 2 separate timescales. (c) Distributions of the number of elementary rearrangements along the shortest pathway from each local minimum to the global minimum for *U-mA, GP-mA*, and *G-mA* models. The number of elementary steps corresponds to the minimum number of transition-state crossings required to reach the global minimum from a given local minimum.

The probability distributions of ln(*τ*_MFPT_) for the Chain B (Figure 8a) models exhibit distinct kinetic characteristics that reflect the underlying organization of their free-energy landscapes. The *U-mB* and *GP-mB* models both exhibit unimodal relaxation-time distributions, with glycosylation broadening the distribution while retaining a single dominant timescale, indicating increased kinetic heterogeneity and sampling of a diverse range of metastable conformations. In contrast to the *G-mA* model, the glycosylated *G-mB* model exhibits a remarkably narrow unimodal distribution. This observation indicates that, although glycosylation increases the structural complexity of the free-energy landscape, relaxation to the global minimum remains dominated by a single characteristic kinetic process. This interpretation is further supported by the distribution of the number of elementary rearrangements, where most local minima require only 2 elementary steps to reach the global minimum, with only a few minima requiring 3 steps as shown in Figure 8c.

**FIG. 8.**
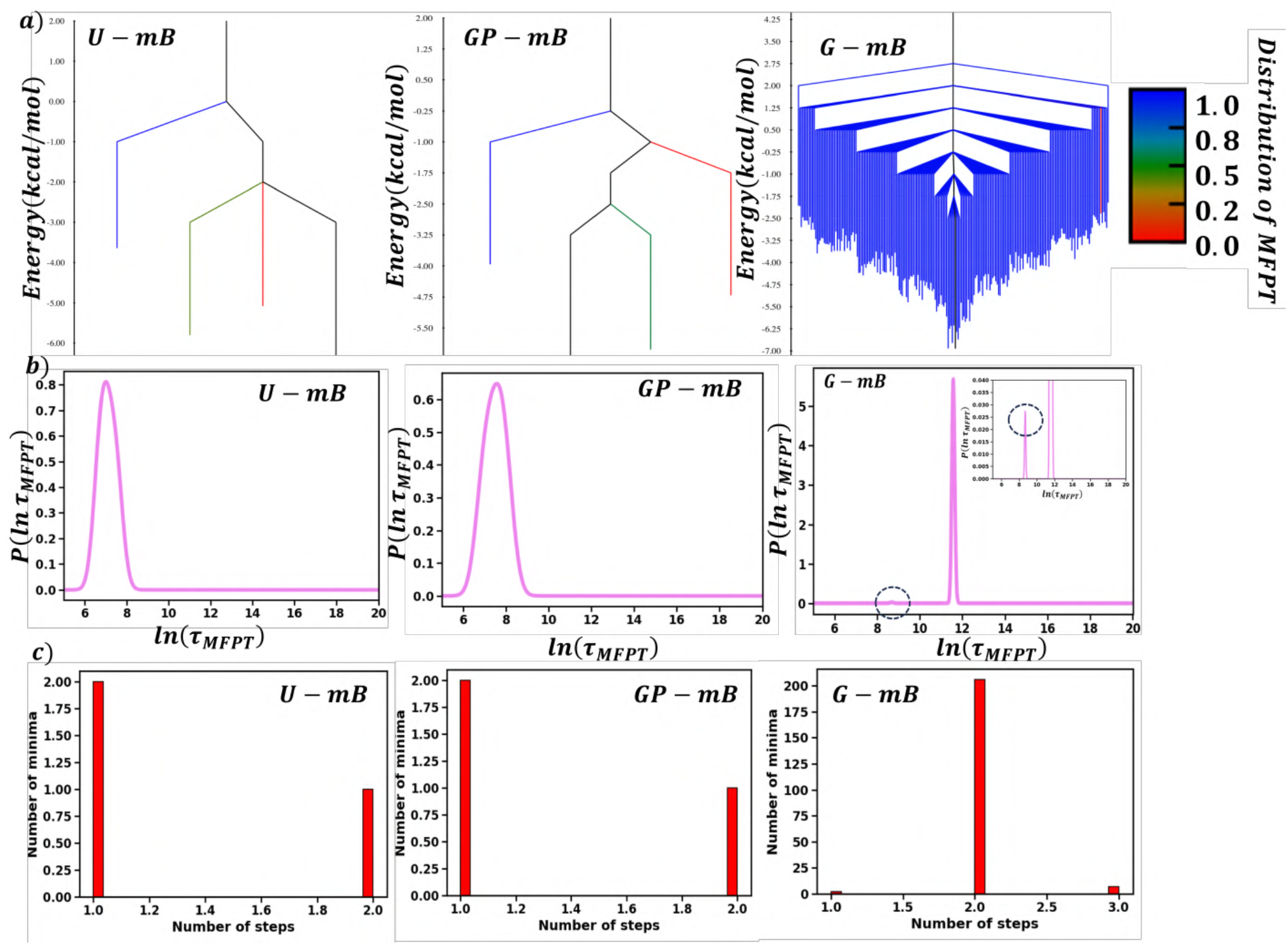
(a) Disconnectivity graphs and (b) probability distributions of the logarithm of the mean first-passage time, *P*(ln *τ*_MFPT_), for *U-mB, GP-mB*, and *G-mB* models. Panel (a) presents the disconnectivity graphs constructed from the local minima. The branches are colored according to the normalized ln *τ*_MFPT_ distribution associated with the corresponding minima, as indicated by the color bar. Panel (b) shows the probability distributions of ln(*τ*_MFPT_) calculated from the MFPT values of all local minima with respect to the global minimum. The distributions characterize the kinetic timescales governing relaxation over the respective energy landscapes. (c) Distributions of the number of elementary rearrangements along the shortest pathway from each local minimum to the global minimum for *U-mB, GP-mB*, and *G-mB* models. The number of elementary steps corresponds to the minimum number of transition-state crossings required to reach the global minimum from a given local minimum.

The probability distributions of ln(*τ*_MFPT_) for the dimeric models reveal distinct kinetic signatures associated with increasing structural complexity upon glycosylation (Figure 9(a)).The *U-dAB* model exhibits a narrow unimodal distribution, indicating relaxation toward the global minimum through a single dominant kinetic process. In contrast, the *GP-dAB* model displays a narrow distribution with a shoulder at longer relaxation times, reflecting the emergence of slower relaxation pathways and increased kinetic heterogeneity induced by glycosylation. The *G-dAB* model exhibits a dominant narrow peak at shorter relaxation times accompanied by weak secondary features extending toward longer timescales. This distribution indicates that the majority of local minima rapidly relax to the global minimum through a common kinetic pathway. The minor peaks correspond to longer relaxation timescales associated with a small population of local minima (Figure 9a,b). The distribution of elementary rearrangements along the MFEPs leading to the global minimum further supports our interpretation. As shown in Figure 9c, nearly all local minima reach the global minimum in a single elementary step, and only a few require 2 steps, highlighting the efficient kinetic connectivity of the dimeric glycosylated landscape despite its increased structural complexity.

**FIG. 9.**
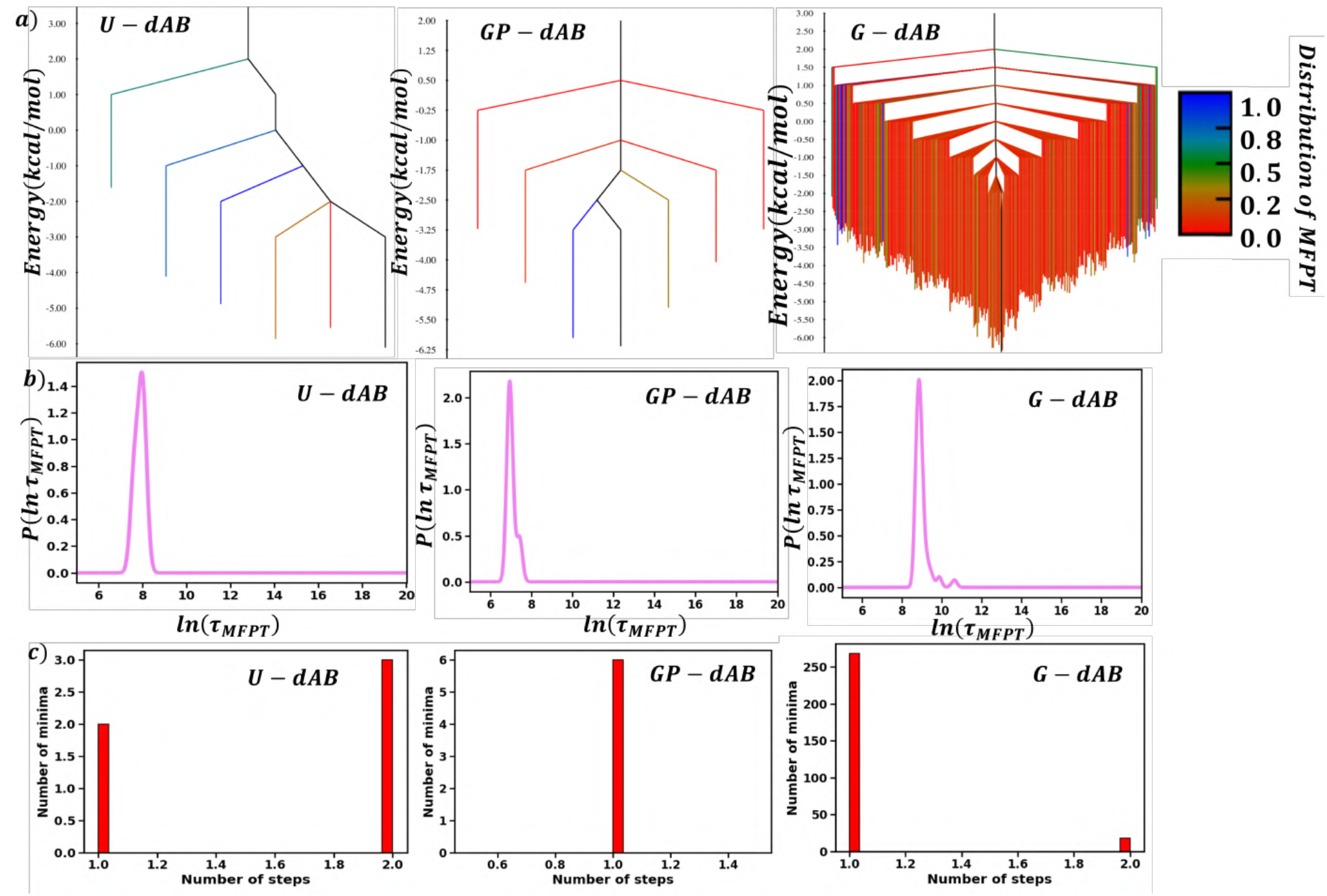
(a) Disconnectivity graphs and (b) probability distributions of the logarithm of the mean first-passage time, *P*(ln *τ*_MFPT_), for *U-dAB, GP-dAB*, and *G-dAB* models. Panel (a) presents the disconnectivity graphs constructed from the local minima. The branches are colored according to the normalized ln *τ*_MFPT_ distribution associated with the corresponding minima, as indicated by the color bar. Panel (b) shows the probability distributions of ln(*τ*_MFPT_) calculated from the MFPT values of all local minima with respect to the global minimum. The distributions characterize the kinetic timescales governing relaxation over the respective energy landscapes. (c) Distributions of the number of elementary rearrangements along the shortest pathway from each local minimum to the global minimum for *U-dAB, GP-dAB*, and *G-dAB* models. The number of elementary steps corresponds to the minimum number of transition-state crossings required to reach the global minimum from a given local minimum.

### H. Quantifying landscape complexity: Shannon entropy and frustration metric

The conformational diversity of the free-energy landscape was quantified using the Shannon entropy, *s*(*T*), which provides a thermodynamic measure of the distribution of equilibrium populations among the free-energy minima. Crucially, the Shannon entropy depends solely on the equilibrium occupation probabilities of the minima and therefore does not require information regarding the transition states or kinetic connectivity of the landscape.

The Shannon entropy is defined as^80^

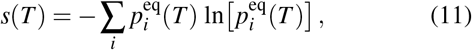

where 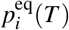 denotes the equilibrium occupation probability of free energy minimum *i* at temperature *T*. Larger values of *s*(*T*) correspond to a broader distribution of populated minima, indicating greater conformational heterogeneity, whereas smaller values imply that the equilibrium population is dominated by a few local minima.

Figure 10 illustrates the temperature dependence of the Shannon entropy for the monomeric and dimeric HCA IX-c systems. As expected, the entropy increases monotonically with temperature as thermal activation progressively populates higher free energy minima, resulting in a broader distribution of accessible conformational states.

**FIG. 10.**
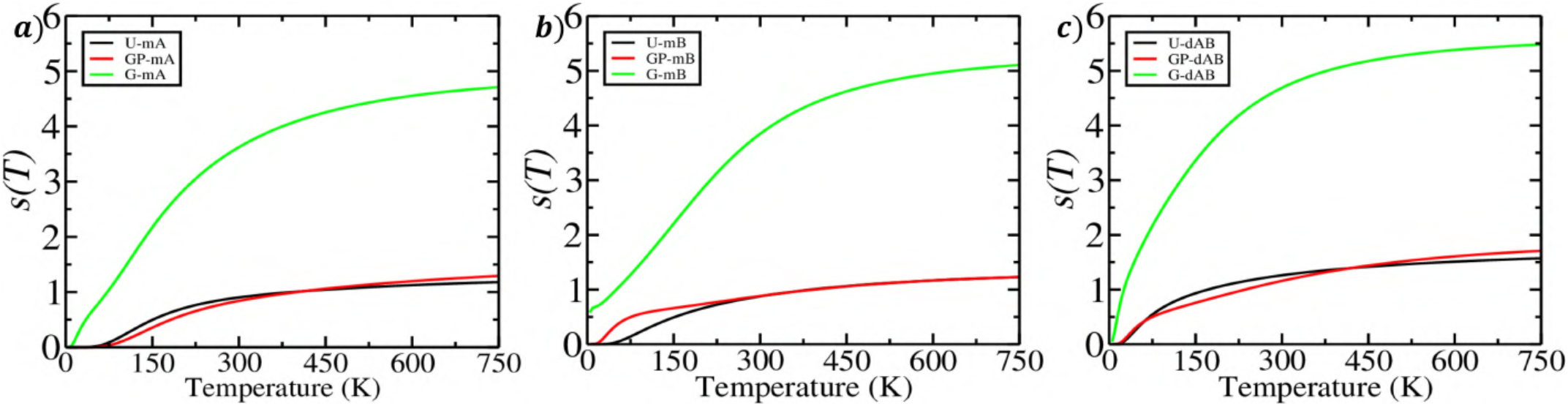
Temperature dependence of the Shannon entropy, *s(T)*, obtained from the free energy disconnectivity graphs (FEDGs) for (a) Chain A, (b) Chain B, and (c) the dimeric HCA IX-c. Black, red, and green curves correspond to the un-glycosylated protein (U), the protein component of the glycoprotein (GP), and the complete glycoprotein (G), respectively. The increase in *s(T)* with temperature reflects the progressive population of higher-energy metastable basins as free energy barriers are overcome. Comparison of the *U* and *GP* curves quantifies the effect of glycosylation on the conformational landscape of the protein alone, whereas the *G* curves include the combined conformational contributions of both the protein and glycan chains. The substantially larger entropy of the complete glycoprotein reflects the enhanced configurational diversity and hierarchical complexity introduced by glycan dynamics.

The un-glycosylated models (*U-mA, U-mB*, and *U-dAB*) exhibit the lowest Shannon entropy over the entire temperature range, consistent with the relatively simple funnel-like organization of their FEDGs, where only 4 − 6 low-energy minima contribute significantly to the equilibrium ensemble. The models *GP-mA, GP-mB*, and *GP-dAB* results in a modest increase in entropy, reflecting the stabilization of additional metastable conformations while largely preserving the underlying funnel-shaped topology.

In contrast, the fully glycosylated systems (*G-mA, G-mB*, and *G-dAB*) display substantially larger Shannon entropy throughout the investigated temperature range. The markedly higher entropy indicates that glycosylation substantially broadens the conformational ensemble by stabilizing a significantly larger number of energetically competitive free-energy minima. Among all the systems investigated, the dimeric glycosylated model (*G-dAB*) exhibits the largest Shannon entropy, demonstrating that the combined effects of glycosylation and dimerization produce the greatest conformational heterogeneity and the most structurally diverse free-energy landscape.

The complexity of the free-energy landscape was further quantified using the frustration metric, *f* (*T*), introduced by Wales and coworkers.^81^ This metric characterizes the efficiency with which a system relaxes toward the global free energy minimum by combining thermodynamic occupation probabilities with the barrier heights separating metastable minima from the global minimum. Unlike measures that depend solely on equilibrium populations, the frustration metric explicitly incorporates the connectivity of the landscape and therefore provides a quantitative description of both the thermodynamic and kinetic contributions to landscape complexity.

The frustration metric was evaluated from the connected network of minima and transition states according to Equation 12,^80^ where the contribution from each metastable minimum is weighted by its equilibrium occupation probability and the relative barrier height along the lowest-energy pathway connecting that minimum to the global minimum. To remove the trivial temperature dependence arising from the increasing equilibrium occupation of the global minimum at low temperatures, the renormalized frustration index, 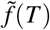, was also calculated using the renormalized equilibrium occupation probabilities, 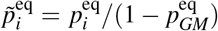.^80^ This renormalization excludes the contribution of the global minimum, allowing direct comparison of the degree of frustration over the entire temperature range.^73^

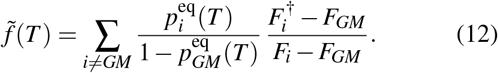

Figure 11 shows the temperature dependence of both *f* (*T*) and 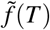 for the monomeric and dimeric HCA IX-c systems. The un-glycosylated models exhibit comparatively low frustration throughout the investigated temperature range, consistent with the relatively simple funnel-like organization of their free energy landscapes. In *protein only* models (GP), there is a moderate increase in frustration, reflecting the stabilization of additional metastable conformational states. This observation is largely consistent with the MFPT distributions presented in the previous section. The fully glycosylated systems exhibit substantially larger values of both *f* (*T*) and 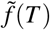, particularly at low temperatures, indicating that glycosylation markedly increases the topological complexity of the free-energy landscape. As the temperature increases, both frustration measures decrease monotonically because thermal activation progressively increases the accessibility of higher-energy pathways, thereby reducing the relative influence of individual kinetic bottlenecks on relaxation toward the global minimum.

**FIG. 11.**
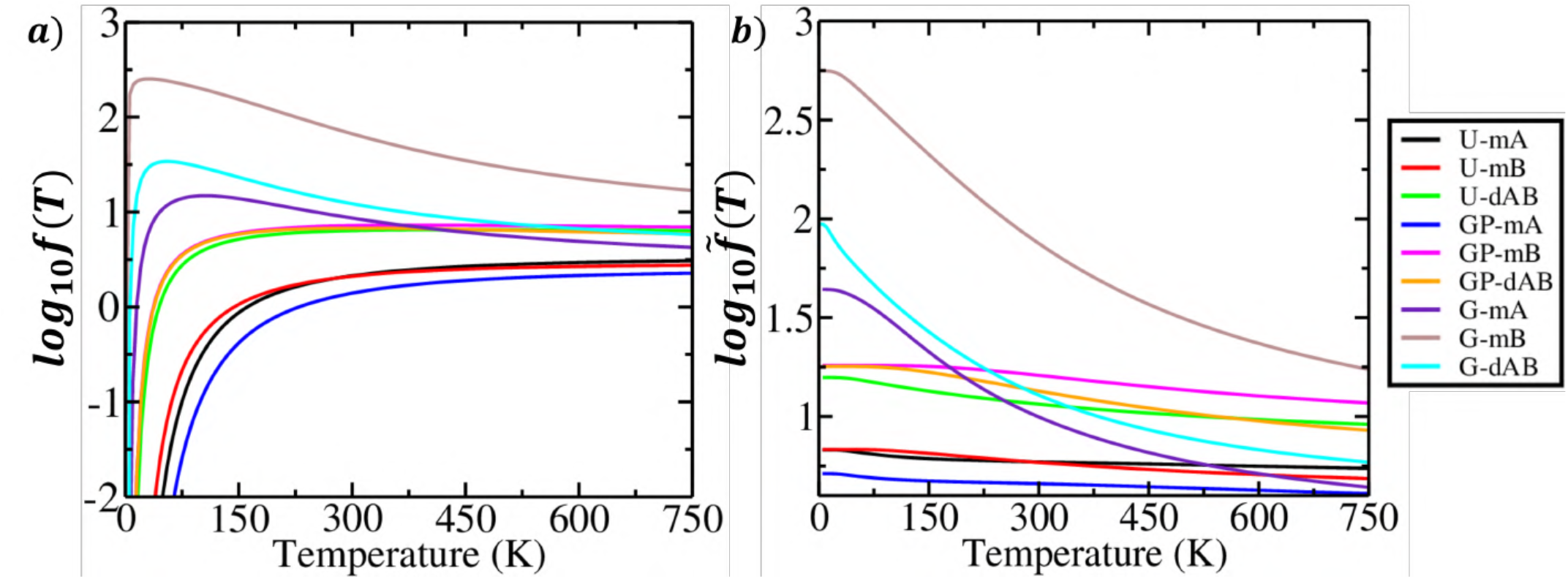
The logarithmic value of (a) frustration index, *f(T)* and (b) normalised frustration index, 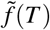 computed as function of temperature from the transition network for all the models.

### I. Free-Energy Disconnectivity Graphs Along the Slow Dynamical Components

To establish the connection between the free-energy landscapes and the underlying slow collective dynamics, the FEDGs were colored according to the first two time-independent components, TIC1 and TIC2, obtained by projecting the high-dimensional CV space comprising glycan dihedral angles and glycan-protein heavy atom distance descriptors onto the tICA space. For *G-mA* (Figures 12a,b), TIC1 provides a clear dynamical distinction between the two funnels: the funnel containing the global minimum is predominantly associated with a higher normalized TIC1 values, whereas the second funnel is characterized by lower TIC1 values. This separation suggests that TIC1 captures the slow collective motion associated with inter-funnel transitions. In contrast, TIC2 varies more continuously across both funnels, indicating that it primarily describes conformational fluctuations within the individual basins. For *G-mB* (Figures 12c,d), which exhibits a single-funnel topology, both components vary continuously across the landscape, with TIC1 showing the dominant gradient along the funnel and TIC2 contributing mainly to local conformational heterogeneity. Similarly, *G-dAB* (Figures 12e-h) exhibits broadly distributed color gradients rather than discrete dynamical partitions. The more pronounced variation in TIC1 indicates its greater contribution to the dominant slow dynamics, whereas TIC2 captures secondary fluctuations within the interconnected metastable network. Overall, tICA provides the clearest dynamical separation when there is a separation of time-scales between local equilibration and inter-basin transitions. Even in such scenarios, tICA may not be a perfect ‘reaction coordinate’, as implied by the intermixing of colors in the FEDG. Thus, lowdimensional projections must be interpreted with caution, especially when the dynamics is characterized by time-scales exhibiting little or no spectral gap.

**FIG. 12.**
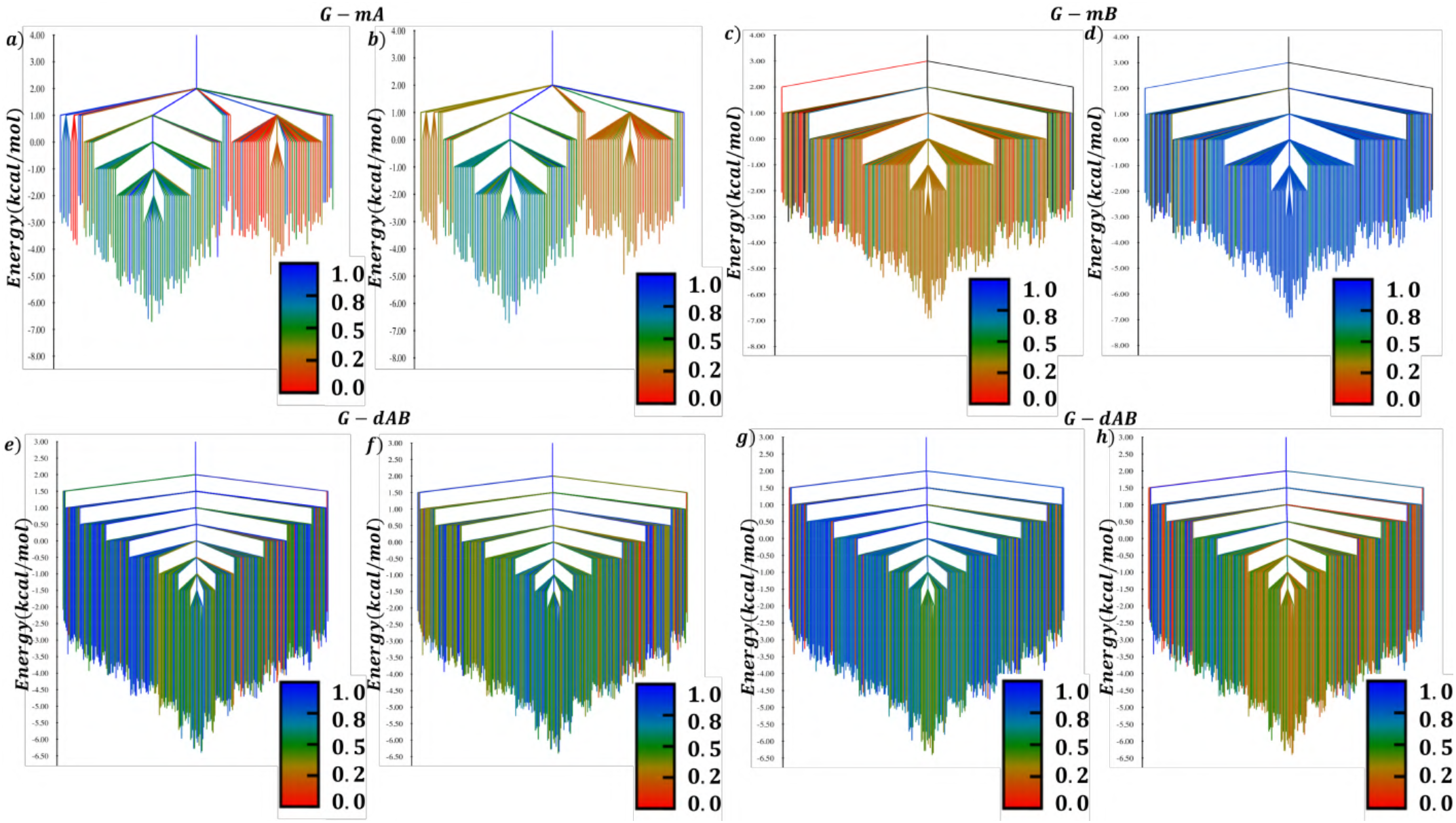
FEDGs coloured according to the projections onto the first two time-structure independent components obtained from tICA analysis. The colour scale represents the normalized values of TIC1 for panels (a), (c), (e), and (g), and TIC2 for panels (b), (d), (f), and (h), highlighting the slow collective conformational dynamics across the free-energy landscapes. Panels (a,b) correspond to *G-mA*, (c,d) to *G-mB*, while panels (e,f) and (g,h) show the respective tICA projections obtained from the glycan-protein CVs of chain A and chain B of the dimeric *G-dAB* model.

## IV. CONCLUSION

In the present work, we have investigated the influence of N-linked glycosylation on the free-energy landscape of the catalytic domain of human carbonic anhydrase IX (HCA IXc) in both its monomeric and dimeric forms. By combining extensive all-atom molecular dynamics simulations with kinetic transition network analysis, free-energy dysconnectivity graphs (FEDGs), minimum free-energy pathways (MFEPs), and mean first-passage time (MFPT) calculations, we have obtained a comprehensive description of the thermodynamic and kinetic organization of the conformational landscape.

The analyses demonstrate that the un-glycosylated systems are characterized by relatively simple funnel-shaped free-energy landscapes containing only four low-energy minima for the monomeric systems and six minima for the dimer, indicating limited conformational heterogeneity. Upon glycosylation, the complexity of the landscape increases dramatically, giving rise to 150, 216, and 287 metastable minima for the *G-mA, G-mB*, and *G-dAB* systems, respectively. Despite this substantial increase in landscape ruggedness, the overall tertiary structure and secondary structural architecture of HCA IX-c remain largely preserved, indicating that glycosylation primarily reshapes the underlying free-energy landscape without disrupting the native protein fold.

The disconnectivity graphs further reveal that the intrinsic conformational flexibility of the N-linked glycan, arising from numerous accessible combinations of the glycosidic torsional angles (*φ, ψ*, and *ω*), is the principal source of the observed landscape complexity. Glycosylation partitions the native protein funnel into multiple interconnected low-energy basins, thereby expanding the accessible conformational ensemble. While chain A exhibits a pronounced double-funnel topology, chain B is characterized by a single dominant funnel, demonstrating that identical glycan chains can reorganize the conformational landscapes of the two monomers in distinct ways.

The kinetic analyses further reveal significant differences in the accessibility of glycan conformational states. The MFEP connecting extended and backfolded conformations proceeds through multiple intermediate minima in *G-mA*, a single intermediate minimum in *G-mB*, and two intermediate minima in the dimeric *G-dAB* system. Consistent with these pathway lengths, the calculated transition rates indicate that backfolded conformations are both thermodynamically and kinetically favored in all glycosylated systems. The MFPT analysis likewise demonstrates that glycosylation broadens the distribution of relaxation timescales relative to un-glycosylated systems, reflecting the emergence of additional kinetically distinct metastable states.

Dimerization further amplifies the complexity of the freeenergy landscape by increasing both the number and connectivity of low-energy basins while preserving efficient kinetic communication between the dominant conformational states. Together, the FEDG, MFEP, and MFPT analyses demonstrate that glycosylation modulates the conformational dynamics of HCA IX-c through a hierarchical reorganization of the underlying free-energy landscape rather than through large-scale structural rearrangements of the catalytic domain.

Overall, this work establishes free-energy disconnectivity graphs, in conjunction with kinetic transition network analysis, is a powerful framework for elucidating the thermodynamic and kinetic organization of glycoprotein free-energy landscapes. Beyond HCA IX-c, the methodology presented here should be broadly applicable to other glycoproteins, providing mechanistic insights into how glycan conformational dynamics regulate protein conformational heterogeneity, kinetic accessibility, and biological function.

## SUPPLEMENTARY MATERIAL

The supporting information includes the multidimensional free-energy surfaces projected onto the first two time-lagged independent components (tICA), the complete definitions of the collective variables employed in the Markov state model analysis, VAMP score comparisons, determination of the optimal clustering cutoff, secondary structure and RMSF analyses, and electrostatic interaction energy networks associated with the highest free-energy barriers along the minimum free-energy pathways.

## ACKNOWLEDGMENTS

R.D. thanks MoE, Government of India for the PMRF fellowship. D.M thanks IIT Kharagpur for providing the research fellowship. The research presented in this article is funded by the Science and Engineering Research Board (SERB), India (grant number CRG/2018/ 002005 and CRG/2022/007493) and Anusandhan National Research Foundation (ANRF) (grant number ANRF/ARG/2025/007845/CS). Additional computational support received from the HPC facility created under the DST-FIST scheme (SR/FST/CSII-011/2005 and SR/FST/CSII-026/2013) and that under the National Supercomputing Mission (NSM), Government of India, supported by the Centre for Development of Advanced Computing (CDAC), Pune is gratefully acknowledged.

## References

1 R. Callender and R. B. Dyer, “The dynamical nature of enzymatic catalysis,” Accounts of Chemical Research 48, 407–413 (2015).

2 P. K. Agarwal, “A biophysical perspective on enzyme catalysis,” Biochemistry 58, 438–449 (2019).

3 P. E. Leopold, M. Montal, and J. N. Onuchic, “Protein folding funnels: a kinetic approach to the sequence-structure relationship.” Proceedings of the National Academy of Sciences USA 89, 8721–8725 (1992).

4 H. Frauenfelder, S. G. Sligar, and P. G. Wolynes, “The energy landscapes and motions of proteins,” Science 254, 1598–1603 (1991).

5 J. N. Onuchic and P. G. Wolynes, “Theory of protein folding,” Current Opinion in Structural Biology 14, 70–75 (2004).

6 K. A. Dill and H. S. Chan, “From levinthal to pathways to funnels,” Nature Structural Biology 4, 10–19 (1997).

7 S. Kumar, B. Ma, C.-J. Tsai, N. Sinha, and R. Nussinov, “Folding and binding cascades: Dynamic landscapes and population shifts,” Protein Science9, 10–19 (2000).

8 J. Wang and G. M. Verkhivker, “Energy landscape theory, funnels, specificity, and optimal criterion of biomolecular binding,” Physical Review Letters 90, 188101 (2003).

9 D. J. Wales, “The energy landscape as a unifying theme in molecular science,” Philosophical Transactions of the Royal Society A: Mathematical, Physical and Engineering Sciences 363, 357–377 (2005).

10 S. V. Krivov and M. Karplus, “Free energy disconnectivity graphs: Application to peptide models,” The Journal of Chemical Physics 117, 10894–10903 (2002).

11 D. J. Wales, J. P. Doye, M. A. Miller, P. N. Mortenson, and T. R. Walsh,“Energy landscapes: from clusters to biomolecules,” Advances in Chemical Physics 115, 1–111 (2000).

12 D. J. Wales and T. V. Bogdan, “Potential energy and free energy landscapes,” (2006).

13 S. C. Frost, Carbonic Anhydrase: Mechanism, Regulation, Links to Disease, and Industrial Applications (Springer Netherlands, 2014) pp. 9–30.

14 A. Scozzafava and C. T. Supuran, “Glaucoma and the applications of carbonic anhydrase inhibitors,” in Carbonic Anhydrase: Mechanism, Regulation, Links to Disease, and Industrial Applications, edited by S. C. Frost and R. McKenna (Springer Netherlands, 2014) pp. 349–359.

15 W. Sly, D. Hewett-Emmett, M. Whyte, Y. Yu, and R. Tashian, “Carbonic anhydrase ii deficiency identified as the primary defect in the autosomal recessive syndrome of osteopetrosis with renal tubular acidosis and cerebral calcification,” Proceedings of the National Academy of Sciences USA 80, 2752–2756 (1983).

16 H. Soda, S. Yukizane, I. Yoshida, S. Aramaki, and H. Kato, “Carbonic anhydrase ii deficiency in a japanese patient produced by a nonsense mutation at tyr-40 in exon 2,” Human Mutation 5, 348–350 (1995).

17 D. Matulis, Carbonic Anhydrase as Drug Target: Thermodynamics and Structure of Inhibitor Binding (Springer International Publishing, 2019).

18 C. Supuran and A. Nocentini, Carbonic Anhydrases: Biochemistry and Pharmacology of an Evergreen Pharmaceutical Target (Elsevier Science, 2019).

19 S. Frost and R. McKenna, Carbonic Anhydrase: Mechanism, Regulation,Links to Disease, and Industrial Applications, Subcellular Biochemistry (Springer Netherlands, 2013).

20 V. Alterio, M. Hilvo, A. Di Fiore, C. T. Supuran, P. Pan, S. Parkkila,A. Scaloni, J. Pastorek, S. Pastorekova, C. Pedone, et al., “Crystal structure of the catalytic domain of the tumor-associated human carbonic anhydrase ix,” Proceedings of the National Academy of Sciences USA 106, 16233–16238 (2009).

21 C. T. Supuran and J.-Y. Winum, “Carbonic anhydrase ix inhibitors in cancer therapy: an update,” Future Medicinal Chemistry 7, 1407–1414 (2015).

22 C. T. Supuran and A. Nocentini, Carbonic anhydrases: biochemistry and pharmacology of an evergreen pharmaceutical target (Academic Press, 2019).

23 S. C. Frost and R. McKenna, Carbonic anhydrase: mechanism, regulation, links to disease, and industrial applications, Vol. 75 (Springer Science & Business Media, 2013).

24 M. Hilvo, L. Baranauskiene, A. M. Salzano, A. Scaloni, D. Matulis, A. Innocenti, A. Scozzafava, S. M. Monti, A. Di Fiore, G. De Simone, et al., “Biochemical characterization of ca ix, one of the most active carbonic anhydrase isozymes,” Journal of Biological Chemistry 283, 27799–27809 (2008).

25 C. C. Wykoff, N. J. Beasley, P. H. Watson, K. J. Turner, J. Pastorek, A. Sibtain, G. D. Wilson, H. Turley, K. L. Talks, P. H. Maxwell, et al., “Hypoxia-inducible expression of tumor-associated carbonic anhydrases,” Cancer Research 60, 7075–7083 (2000).

26 V. Pontecorvi, M. Mori, F. Picarazzi, S. Zara, S. Carradori, A. Cataldi,A. Angeli, E. Berrino, P. Chimenti, A. Ciogli, D. Secci, P. Guglielmi, and C. T. Supuran, “Novel insights on human carbonic anhydrase inhibitors based on coumalic acid: Design, synthesis, molecular modeling investigation, and biological studies,” International Journal of Molecular Sciences 23, 7950 (2022).

27 A. Bonardi and C. T. Supuran, “Polypharmacology of carbonic anhydrase inhibitors and activators,” Expert Opinion on Pharmacotherapy 26, 567–580 (2025).

28 D. Tsikas, “Acetazolamide and human carbonic anhydrases: retrospect, review and discussion of an intimate relationship,” Journal of Enzyme Inhibition and Medicinal Chemistry 39 (2024).

29 D. C. Singleton, A. Macann, and W. R. Wilson, “Therapeutic targeting of the hypoxic tumour microenvironment,” Nature Reviews Clinical Oncology 18, 751–772 (2021).

30 R. Dey and S. Taraphder, “Molecular modeling of glycosylated catalytic domain of human carbonic anhydrase ix,” The Journal of Physical Chemistry B 128, 11054–11068 (2024).

31 R. Dey, K. Shukla, A. Das, and S. Taraphder, “Effect of glycosylation on the reorganization at the active site of human carbonic anhydrase ix,” ChemPhysChem 26, e202500573 (2025).

32 R. Dey and S. Taraphder, “Molecular modeling of glycosylated dimeric catalytic domain of human carbonic anhydrase ix,” (2026), unpublished manuscript.

33 O. M. Becker and M. Karplus, “The topology of multidimensional potential energy surfaces: Theory and application to peptide structure and kinetics,” The Journal of Chemical Physics 106, 1495–1517 (1997).

34 G. J. Rylance, R. L. Johnston, Y. Matsunaga, C.-B. Li, A. Baba, and T. Komatsuzaki, “Topographical complexity of multidimensional energy landscapes,” Proceedings of the National Academy of Sciences USA 103, 18551–18555 (2006).

35 J. A. Joseph, K. Röder, D. Chakraborty, R. G. Mantell, and D. J. Wales, “Exploring biomolecular energy landscapes,” Chemical Communications 53, 6974–6988 (2017).

36 D. Wales, Energy Landscapes: Applications to Clusters, Biomolecules and Glasses, Cambridge Molecular Science (Cambridge University Press, 2004).

37 S. A. M. Stein, A. E. Loccisano, S. M. Firestine, and J. D. Evanseck, “Principal components analysis: a review of its application on molecular dynamics data,” Annual Reports in Computational Chemistry 2, 233–261 (2006).

38 C. C. David and D. J. Jacobs, “Principal component analysis: a method for determining the essential dynamics of proteins,” in Protein dynamics: Methods and Protocols (Springer, 2013) pp. 193–226.

39 A. Altis, P. H. Nguyen, R. Hegger, and G. Stock, “Dihedral angle principal component analysis of molecular dynamics simulations,” The Journal of Chemical Physics 126, 244111 (2007).

40 S. Ma and Y. Dai, “Principal component analysis based methods in bioinformatics studies,” Briefings in Bioinformatics 12, 714–722 (2011).

41 G. Pérez-Hernández, F. Paul, T. Giorgino, G. De Fabritiis, and F. Noé, “Identification of slow molecular order parameters for markov model construction,” Journal of Chemical Physics 139, 015102 (2013).

42 M. M. Sultan and V. S. Pande, “tica-metadynamics: accelerating metady-namics by using kinetically selected collective variables,” Journal of Chemical Theory and Computation 13, 2440–2447 (2017).

43 L. McInnes, J. Healy, N. Saul, and L. Großberger, “Umap: Uniform manifold approximation and projection,” Journal of Open Source Software 3, 861 (2018).

44 V. Spiwok and P. Kriz, “Time-lagged t-distributed stochastic neighbor embedding (t-sne) of molecular simulation trajectories,” Frontiers in Molecular Biosciences 7, 132 (2021).

45 S. V. Krivov and M. Karplus, “Hidden complexity of free energy surfaces for peptide (protein) folding,” Proceedings of the National Academy of Sciences USA 101, 14766–14770 (2004).

46 D. Wales, “Perspective: Insight into reaction coordinates and dynamics from the potential energy landscape,” The Journal of Chemical Physics 142(2015).

47 L. Bonati, G. Piccini, and M. Parinello, “Deep learning the slow modes for rare events sampling,” Proceedings of the National Academy of Sciences USA 118, e2113533118 (2024).

48 D. J. Wales, M. A. Miller, and T. R. Walsh, “Archetypal energy landscapes,”Nature 394, 758–760 (1998).

49 L. C. Smeeton, M. T. Oakley, and R. L. Johnston, “Visualizing energy landscapes with metric disconnectivity graphs,” Journal of Computational Chemistry 35, 1481–1490 (2014).

50 J. P. K. Doye, M. A. Miller, and D. J. Wales, “The double-funnel energy landscape of the 38-atom lennard-jones cluster,” The Journal of Chemical Physics 110, 6896 (1999).

51 T. James, D. J. Wales, and J. H. Rojas, “Energy landscapes of water clusters in a uniform electric field,” The Journal of Chemical Physics 126, 054506 (2007).

52 D. A. Evans and D. J. Wales, “Free energy landscapes of model peptides and proteins,” The Journal of Chemical Physics 118, 3891 (2003).

53 D. Chakraborty, R. Collepardo-Guevara, and D. J. Wales, “Energy landscapes, folding mechanisms, and kinetics of rna tetraloop hairpins,” Journalof the American Chemical Society 136, 18052–18061 (2014).

54 S. Niblett, V. K. de Souza, J. D. Stevenson, and D. J. Wales, “Dynamics of a molecular glass former: Energy landscapes for diffusion in orthoterphenyl,” The Journal of Chemical Physics 145, 024505 (2016).

55 C. Tian, K. Kasavajhala, K. A. A. Belfon, L. Raguette, H. Huang, A. N. Migues, J. Bickel, Y. Wang, J. Pincay, Q. Wu, and C. Simmerling, “ff19sb: Amino-acid-specific protein backbone parameters trained against quantum mechanics energy surfaces in solution,” Journal of Chemical Theory and Computation 16, 528–552 (2020).

56 M. Basma, S. Sundara, D. Çalgan, T. Vernali, and R. J. Woods, “Solvated ensemble averaging in the calculation of partial atomic charges,” Journal of Computational Chemistry 22, 1125–1137 (2001).

57 A. Sarkar and S. Pérez, “Protein–carbohydrate interactions,” Structural Glycobiology, 71 (2012).

58 R. J. Woods, R. A. Dwek, C. J. Edge, and B. Fraser-Reid, “Molecular mechanical and molecular dynamic simulations of glycoproteins and oligosaccharides. 1. glycam_93 parameter development,” The Journal of Physical Chemistry 99, 3832–3846 (1995).

59 R. Woods and R. Chappelle, “Restrained electrostatic potential atomic partial charges for condensed-phase simulations of carbohydrates,” Journal of Molecular Structure: THEOCHEM 527, 149–156 (2000).

60 M. B. Peters, Y. Yang, B. Wang, L. Fusti-Molnar, M. N. Weaver, and K. M. Merz Jr, “Structural survey of zinc-containing proteins and development of the zinc amber force field (zaff),” Journal of Chemical Theory and Computation 6, 2935–2947 (2010).

61 P. Mark and L. Nilsson, “Structure and dynamics of the tip3p, spc, and spc/e water models at 298 k,” The Journal of Physical Chemistry A 105, 9954–9960 (2001).

62 T. Zhou and A. Caflisch, “Distribution of reciprocal of interatomic distances: A fast structural metric,” Journal of Chemical Theory and Computation 8, 2930–2937 (2012).

63 M. Schaffler, D. J. Wales, and B. Strodel, “Energy landscape and kinetic analysis of molecular dynamics simulations for intrinsically disordered proteins,” The Journal of Physical Chemistry B 129, 11430–11440 (2025).

64 M. K. Scherer, B. Trendelkamp-Schroer, F. Paul, G. Perez-Hernandez,M. Hoffmann, N. Plattner, C. Wehmeyer, J.-H. Prinz, and F. Noe, “Pyemma 2: A software package for estimation, validation, and analysis of markov models,” Journal of Chemical Theory and Computation 11, 5525–5542 (2015).

65 I. Y. Ibrahim and I. M. Zebari, “A comprehensive review of the fordfulkerson algorithm for network flow problems,” Asian Journal of Research in Computer Science 18, 335–344 (2025).

66 R. E. Gomory and T. C. Hu, “Multi-terminal network flows,” Journal of the Society for Industrial and Applied Mathematics 9, 551 (1961).

67 A. A. Hagberg, D. A. Schult, and P. J. Swart, “Exploring network structure, dynamics, and function using networkx,” in Proceedings of the 7th Python in Science Conference (SciPy 2008), edited by G. Varoquaux, T. Vaught, and J. Millman (Pasadena, CA USA, 2008) pp. 11–15.

68 E. L. Platt, Network science with Python and NetworkX quick start guide:explore and visualize network data effectively (Packt Publishing Ltd, 2019).

69 D. J. Wales, “PATHSAMPLE: A program for generating connected stationary-point databases and extracting global kinetics,” (2003), accessed: 2026-08-06.

70 E. W. Dijkstra, “A note on two problems in connexion with graphs,” Numerische Mathematik, 269–271 (1959).

71 D. J. Wales, “Calculating rate constants and committor probabilities fortransition networks by graph transformation,” The Journal of Chemical Physics 130 (2009).

72 M. A. Miller, D. J. Wales, and V. K. de Souza, “disconnectionDPS,” (1998).

73 D. Chakraborty, J. E. Straub, and D. Thirumalai, “Energy landscapes of aβ monomers are sculpted in accordance with ostwald’s rule of stages,” Science Advances 9, eadd6921 (2023).

74 M. Schaffler, B. Strodel, and D. J. Wales, “The energy landscape of aβ 42: a funnel to disorder for the monomer becomes a folding funnel for self-assembly,” Chemical Communications 60, 13574–135577 (2024).

75 D. Chakraborty, Y. Chebaro, and D. J. Wales, “A multifunnel energy landscape encodes the competing α-helix and β-hairpin conformations for a designed peptide,” Physical Chemistry Chemical Physics 22, 1359–1370 (2020).

76 D. Chakraborty and D. J. Wales, “Dynamics of an adenine-adenine rna conformational switch from discrete path sampling,” The Journal of Chemical Physics 150, 125101 (2019).

77 D. Chakraborty and D. J. Wales, “Energy landscape and pathways for transitions between watson-crick and hoogsteen base-pairing in dna,” The Journal of Physical Chemistry Letters 9, 229–241 (2018).

78 D. J. Wales, “Calculating rate constants from networks of transition states,”The Journal of Chemical Physics 124, 234110 (2006).

79 D. J. Wales, “Dynamical signatures of multifunnel energy landscapes,” The Journal of Physical Chemistry Letters 13, 6349 (2022).

80 V. De Souza, J. Stevenson, S. Niblett, J. Farrell, and D. Wales, “Defining and quantifying frustration in the energy landscape: Applications to atomic and molecular clusters, biomolecules, jammed and glassy systems,” The Journal of Chemical Physics 146 (2017).

81 V. K. de Souza, J. D. Stevenson, S. P. Niblett, J. D. Farrell, and D. J. Wales, “Defining and quantifying frustration in the energy landscape: Applications to atomic and molecular clusters, biomolecules, jammed and glassy systems,” The Journal of Chemical Physics 146, 124103 (2017).

82 F. H. Stillinger and T. A. Weber, “Packing structures and transitions in liquids and solids,” Science 225, 983–989 (1984).

83 A. R. Oganov and M. Valle, “How to quantify energy landscapes of solids,” The Journal of Chemical Physics 130 (2009).

84 J. P. Doye and C. P. Massen, “Characterizing the network topology of the energy landscapes of atomic clusters,” The Journal of Chemical Physics122 (2005).

85 B. W. Shires and C. J. Pickard, “Visualizing energy landscapes through manifold learning,” Physical Review X 11, 041026 (2021).

86 S. V. Krivov and M. Karplus, “One-dimensional free-energy profiles of complex systems: Progress variables that preserve the barriers,” The Journal of Physical Chemistry B 110, 12689–12698 (2006).

87 S. V. Krivov and M. Karplus, “Diffusive reaction dynamics on invariant free energy profiles,” Proceedings of the National Academy of Sciences USA 105, 13841–13846 (2008).

88 F. Noé and S. Fischer, “Transition networks for modeling the kinetics of conformational change in macromolecules,” Current Opinion in Structural Biology 18, 154–162 (2008).

89 D. J. Wales, “Energy landscapes: calculating pathways and rates,” International Reviews in Physical Chemistry 25, 237–282 (2006).

90 S. Prasad, F. Aviat, I. Gonzales, James E., and B. R. Brooks, “apocharmm: High-performance molecular dynamics simulations on gpus for advanced simulation methods,” The Journal of Chemical Physics 162, 182501 (2025).

91 M. Heinig and D. Frishman, “Stride: a web server for secondary structure assignment from known atomic coordinates of proteins,” Nucleic Acids Research 32, 500–502 (2004).

92 S.-J. Yang, D. J. Wales, E. J. Woods, and G. R. Fleming, “Design principles for energy transfer in the photosystem ii supercomplex from kinetic transition networks,” Nature Communications 15, 8763 (2024).

93 J. Cohen, L. A. Rodrigues, and E. P. Duarte Jr., “A parallel implementation of gomory-hu’s cut tree algorithm,” in 2012 IEEE 24th International Symposium on Computer Architecture and High Performance Computing (2012) pp. 124–131.

94 J.-H. Prinz, H. Wu, M. Sarich, B. Keller, M. Senne, M. Held, J. D. Chodera,C. Schütte, and F. Noé, “Markov models of molecular kinetics: Generation and validation.” The Journal of Chemical Physics 134, 174105 (2011).

95 N. F. Polizzi, M. J. Therien, and D. N. Beratan, “Mean first-passage times in biology,” Israel Journal of Chemistry 56, 816–824 (2016).

96 D. J. Sharpe and D. J. Wales, “Graph transformation and shortest paths algorithms for finite markov chains,” Physical Review E 103, 063306 (2021).

97 P. A. Wesołowski, “Multiscale frameworks for exploring protein energy landscapes: advances in theory and simulation: Pa wesołowski,” Journalof Biological Physics 52, 24 (2026).

98 L. Ford and D. R. Fulkerson, “Maximal flow through a network,” Canadian Journal of Mathematics 8, 399 (1956).

