## Supplementary material for "Effect of Glycosylation on the Free Energy Landscape of the Catalytic Domain of Human Carbonic Anhydrase IX": Figure S1

### LIST OF FIGURES

S1 Multi-dimensional free-energy surfaces projected onto the first two time-lagged independent components (IC1 and IC2). Panels (a) and (b) correspond to the glycosylated monomeric model *G-mA* obtained from classical molecular dynamics (MD) and accelerated molecular dynamics (aMD) simulations, respectively of HCA IX-c in water. Panels (c), (d) and (e) show the corresponding free-energy surfaces for the chains *G-mB*, *dA* and *dB*, in glycosylated dimeric HCA IX-c (*G-dAB*). Panels (a) and (b) are reproduced with permission from Ref.[3]. Copyright 2024 American Chemical Society. . . . . 6

S2 Comparison of the VAMP score for different feature representations- DRID (blue), glycan dihedral angles with pair distances (orange), and pair distances only (green), evaluated at lag times of (a) 5 ps, (b) 10 ps, and (c) 20 ps. The representation with DRID exhibits higher VAMP scores across all lag times. . 7

S3 Determination of the optimal number of clusters using regular space clustering at different distance cutoff for all the systems. The optimal distance cutoff for each model was identified from the knee point of the curve, beyond which the number of clusters changes only marginally. The selected cutoff values are indicated by dashed lines. . . . . . . . . . . . . . . . . . . . . . . . . . . . . . . . . . . . . 7

S4 Panels (a), (b) and (c) present the structural integrity for the monomeric and dimeric HCA IX-c from the 3 $\mu$s MD trajectories. . . . . . . . . . . . . . . . . . . . . . . . . 8
S5 Electrostatic component of linear interaction energy networks between the glycan chain and the surrounding protein residues corresponding to the highest free-energy barrier along the minimum free-energy pathway (MFEP). Panels (a), (b), and (c,d) depict the representative transition states connecting minima 7 $\rightarrow$ 64 in *G-mA*, 216 $\rightarrow$ 1 in *G-mB*, and 167 $\rightarrow$ 8 in *G-dAB*, respectively. . . . . . . . . . . . . . . . . . . . . . . . . . . . . . . . . . . . . . . . . . . . . . . . . . 9

### S1. CONFORMATIONAL STATES OF THE GLYCAN CHAIN

To characterize the conformational dynamics of the N-linked glycan chain, we employed a time-lagged independent component analysis (tICA)[4,5] using a comprehensive set of structurally meaningful collective variables (CVs). The feature space comprised two classes of descriptors: (i) all pair distances $pd_{ij}$, between the i$^{\text{th}}$ atom of the glycan chain and j$^{\text{th}}$ protein atom lying within 5 Å from any atom of the glycan chain and (ii) all 36 ($mA$)/ 72 ($dAB$) dihedral angles of the glycan chain(s).[3] The complete definitions of these CVs are provided in Tables S1 and S2. Table S1 lists all glycan dihedral angles considered for the monomeric and dimeric glycosylated systems, while Table S2 summarizes the residue-specific glycan-protein distance descriptors that capture the local interaction network surrounding the glycosylation site. Together, these variables constitute the feature space used for dimensionality reduction and free-energy landscape construction.

The multidimensional free-energy surfaces obtained by projecting the trajectories onto the first two time-lagged independent components (IC1 and IC2) are presented in Figure S1. The figure compares the conformational landscapes of the glycosylated monomeric and dimeric HCA IX-c systems and highlights the dominant metastable basins sampled during the simulations. For the *G-mA* model, the projections obtained from conventional molecular dynamics (MD) and accelerated molecular dynamics (aMD) are shown separately to illustrate the enhanced configurational sampling achieved by aMD.[3] The remaining panels correspond to the glycan chains of *G-mB and the* the glycosylated dimer (*G-dA*, and *G-dB*), revealing distinct conformational ensembles and representative structural states associated with the major free-energy minima. These low-dimensional projections served as the basis of our earlier analysis of glycan conformational heterogeneity and motivated the subsequent development of the high-resolution FEDG[6,7] framework presented in the main manuscript.

Figure S1 compares the free-energy surfaces obtained from conventional molecular dynamics (MD) and accelerated molecular dynamics (aMD) simulations using the first two time-lagged independent components (IC1 and IC2) as reaction coordinates. The conventional MD simulation of the glycosylated monomeric system (*G-mA*) samples two well-defined free-energy minima corresponding to the half-Backfold (H-Bf) and extend-b gg conformations, indicating limited exploration of the conformational landscape within the accessible simulation timescale.[3] In contrast, the aMD simulation significantly broadens the configurational

|  | Definition |  | Definition |
|---|---|---|---|
| CVs for all the Models | Dihedral Angles of the Glycan Chain | CVs for all the Models | Dihedral Angles of the Glycan Chain |
| $\chi_1$ | Asn-212 ND2 - 4YB(a) C1 - C2 - C3 | $\chi_{19}$ | VMB C3 - C4 - C5 - O5 |
| $\chi_2$ | Asn-212 ND2 - 4YB(a) C1 - O5 - C5 | $\chi_{20}$ | VMB C4 - C5 - O5 - C1 |
| $\chi_3$ | 4YB(a) C1 - C2 - C3 - C4 | $\chi_{21}$ | VMB C2 - C3 - O3 - 0MA(1) C1 |
| $\chi_4$ | 4YB(a) C2 - C3 - C4 - C5 | $\chi_{22}$ | VMB C4 - C3 - O3 - 0MA(1) C1 |
| $\chi_5$ | 4YB(a) C3 - C4 - C5 - O5 | $\chi_{23}$ | 0MA(1) C1 - C2 - C3 - C4 |
| $\chi_6$ | 4YB(a) C4 - C5 - O5 - C1 | $\chi_{24}$ | 0MA(1) C2 - C3 - C4 - C5 |
| $\chi_7$ | 4YB(a) C2 - C3 - C4 - O4 | $\chi_{25}$ | 0MA(1) C3 - C4 -C5 - O5 |
| $\chi_8$ | 4YB(a) C3 - C4 - O4 - 4YB(b) C1 | $\chi_{26}$ | 0MA(1) C4 - C5 - O5 - C1 |
| $\chi_9$ | 4YB(a) C5 - C4 - O4 - 4YB(b) C1 | $\chi_{27}$ | 0MA(1) C5 - O5 - C1 - VMB O3 |
| $\chi_{10}$ | 4YB(b) C1 - C2 - C3 - C4 | $\chi_{28}$ | VMB C1 - O5 - C5 - C6 |
| $\chi_{11}$ | 4YB(b) C2 - C3 - C4 - C5 | $\chi_{29}$ | VMB O5 - C5 - C6 - O6 |
| $\chi_{12}$ | 4YB(b) C3 - C4 - C5 - O5 | $\chi_{30}$ | VMB C5 - C6 - O6 - 0MA(2) |
| $\chi_{13}$ | 4YB(b) C4 - C5 - O5 - C1 | $\chi_{31}$ | 0MA(2) C1 - C2 - C3 - C4 |
| $\chi_{14}$ | 4YB(b) C2 - C3 - C4 - O4 | $\chi_{32}$ | 0MA(2) C2 - C3 - C4 - C5 |
| $\chi_{15}$ | 4YB(b) C3 - C4 - O4 - VMB C1 | $\chi_{33}$ | 0MA(2) C3 - C4 - C5 - O5 |
| $\chi_{16}$ | 4YB(b) C5 - C4 - O4 - VMB C1 | $\chi_{34}$ | 0MA(2) C4 - C5 - O5 - C1 |
| $\chi_{17}$ | VMB C1 - C2 - C3 - C4 | $\chi_{35}$ | 0MA(2) C5 - O5 - C1 - VMB O6 |
| $\chi_{18}$ | VMB C2 - C3 - C4 - C5 | $\chi_{36}$ | 0MA(2) O5 - C1 - VMB O6 - C6 |

TABLE S1: List of all the glycan dihedral angles chosen as order parameters for all the models under investigation.

space sampled by the glycan and reveals additional metastable conformations, including the extend-a gt, backfold (Bf) and half-backfold (H-Bf) states as illustrated in Figure S1(b-e).

| | Definition | | | |
|---|---|---|---|---|
| CVs for all the Models | Distance between any $i$-th heavy glycan atom and any $j$-th protein atom within 5 Å of the glycan chain for *G-mA* | Distance between any $i$-th heavy glycan atom and any $j$-th protein atom within 5 Å of the glycan chain for *G-mB* | Distance between any $i$-th heavy glycan atom and any $j$-th protein atom within 5 Å of the glycan chain for *G-dA* | Distance between any $i$-th heavy glycan atom and any $j$-th protein atom within 5 Å of the glycan chain for *G-dB* |
| $d_1$ | Gln-35 | Gln-35 | Gln-35 | Gln-35 |
| $d_2$ | Val-107 | Glu-108 | Val-107 | Val-107 |
| $d_3$ | Glu-108 | His-110 | Glu-108 | Glu-108 |
| $d_4$ | His-110 | Leu-145 | His-110 | His-110 |
| $d_5$ | Glu-146 | Glu-146 | Glu-146 | Glu-146 |
| $d_6$ | Glu-147 | Glu-147 | Glu-147 | Glu-147 |
| $d_7$ | Pro-149 | Pro-149 | Pro-149 | Pro-149 |
| $d_8$ | Glu-150 | Glu-150 | Glu-150 | Glu-150 |
| $d_9$ | Ser-185 | Asp-186 | Ser-185 | Ser-185 |
| $d_{10}$ | Asp-186 | Arg-189 | Asp-186 | Asp-186 |
| $d_{11}$ | Arg-189 | Phe-211 | Arg-189 | Arg-189 |
| $d_{12}$ | Phe-211 | Asn-212 | Phe-211 | Phe-211 |
| $d_{13}$ | Asn-212 | Gln-213 | Asn-212 | Asn-212 |
| $d_{14}$ | Gln-213 | Thr-214 | Gln-213 | Gln-213 |
| $d_{15}$ | Thr-214 | Met-216 | Thr-214 | Thr-214 |
| $d_{16}$ | Met-216 | Pro-257 | Met-216 | Met-216 |
| $d_{17}$ | Phe-256 | | Phe-256 | Phe-256 |
| $d_{18}$ | Pro-257 | | Pro-257 | |

TABLE S2: List of all paired distances chosen as order parameters for all the models under study.

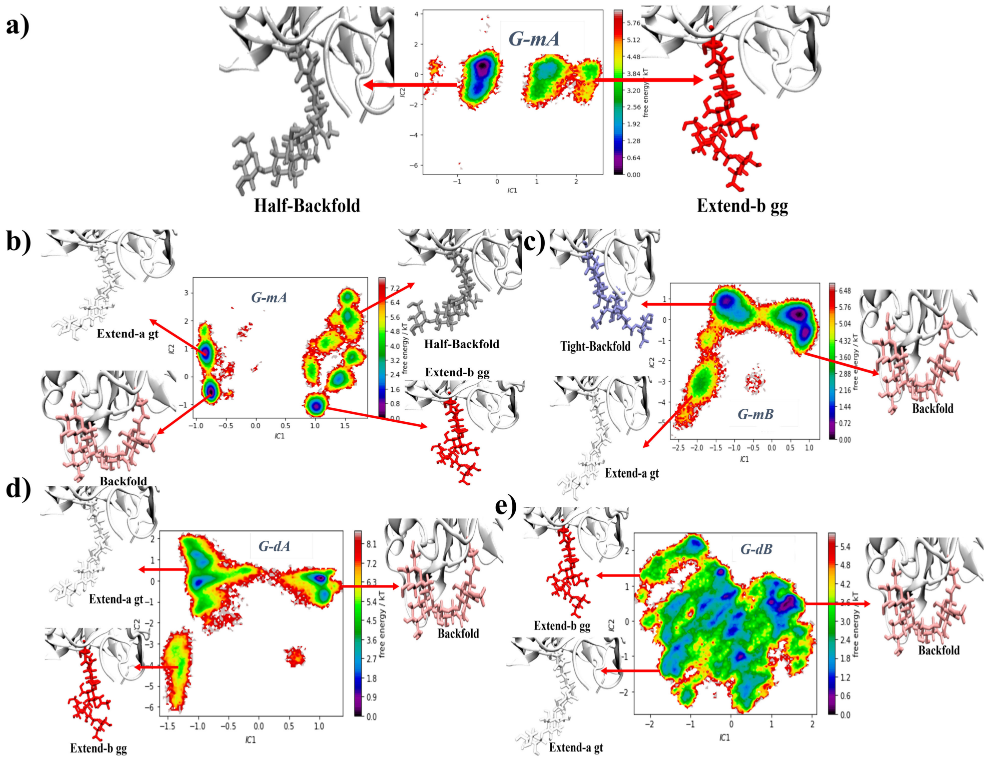

FIG. S1: Multi-dimensional free-energy surfaces projected onto the first two time-lagged independent components (IC1 and IC2). Panels (a) and (b) correspond to the glycosylated monomeric model *G-mA* obtained from classical molecular dynamics (MD) and accelerated molecular dynamics (aMD) simulations, respectively of HCA IX-c in water. Panels (c), (d) and (e) show the corresponding free-energy surfaces for the chains *G-mB*, *dA* and *dB*, in glycosylated dimeric HCA IX-c (*G-dAB*). Panels (a) and (b) are reproduced with permission from Ref.[3]. Copyright 2024 American Chemical Society.

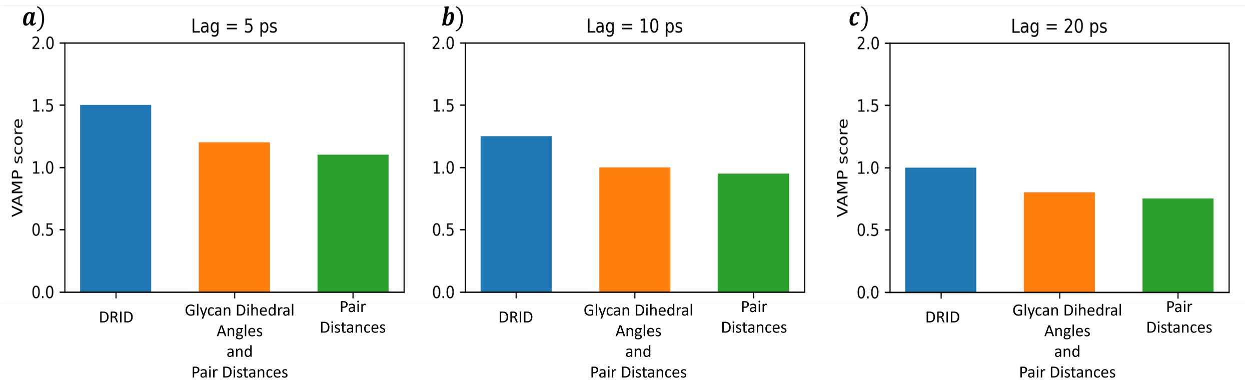

FIG. S2: Comparison of the VAMP score for different feature representations- DRID (blue), glycan dihedral angles with pair distances (orange), and pair distances only (green), evaluated at lag times of (a) 5 ps, (b) 10 ps, and (c) 20 ps. The representation with DRID exhibits higher VAMP scores across all lag times.

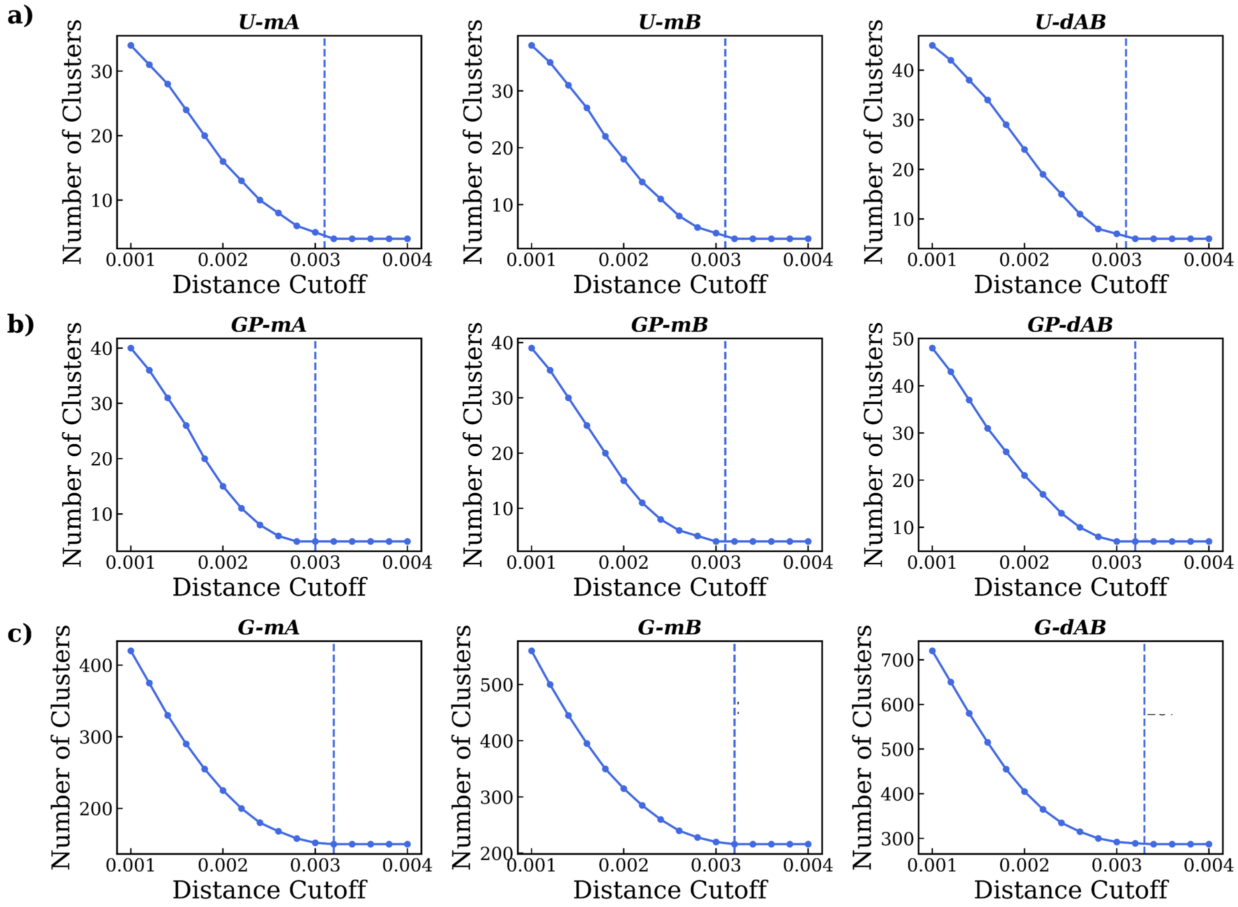

FIG. S3: Determination of the optimal number of clusters using regular space clustering at different distance cutoff for all the systems. The optimal distance cutoff for each model was identified from the knee point of the curve, beyond which the number of clusters changes only marginally. The selected cutoff values are indicated by dashed lines.

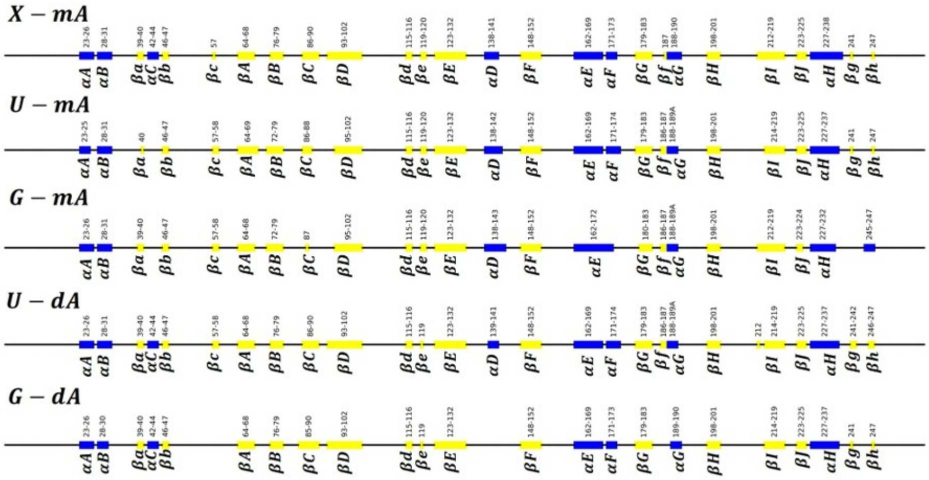

(a) Secondary structure segments obtained from the crystal structure ($X$-$mA$) and at the end of 3 $\mu$s MD trajectories for un-glycosylated (U) and glycosylated (G) monomeric (m) and dimeric (d) chain A of HCA IX-c.

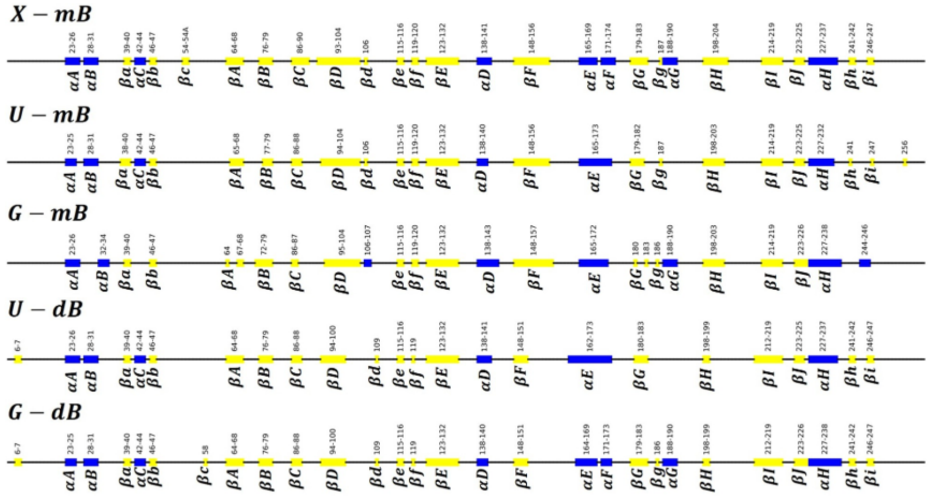

(b) Secondary structure segments obtained from the crystal structure ($X$-$mB$) and at the end of 3 $\mu$s MD trajectories for un-glycosylated (U) and glycosylated (G) monomeric (m) and dimeric (d) chain B of HCA IX-c.

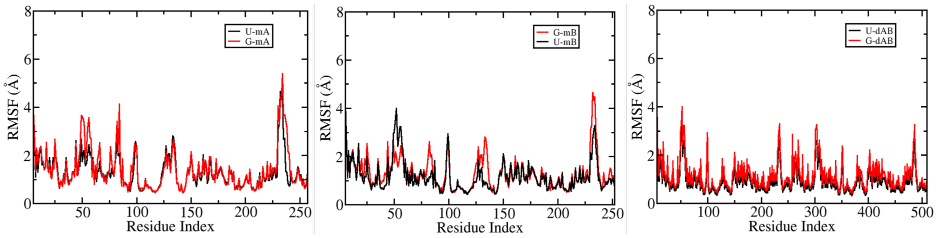

(c) Residue-wise structural fluctuations (RMSF) in the un-glycosylated (U) and glycosylated (G) forms of monomeric (mA, mB) and dimeric (dAB) HCA IX-c in water.

FIG. S4: Panels (a), (b) and (c) present the structural integrity for the monomeric and dimeric HCA IX-c from the 3 $\mu$s MD trajectories.

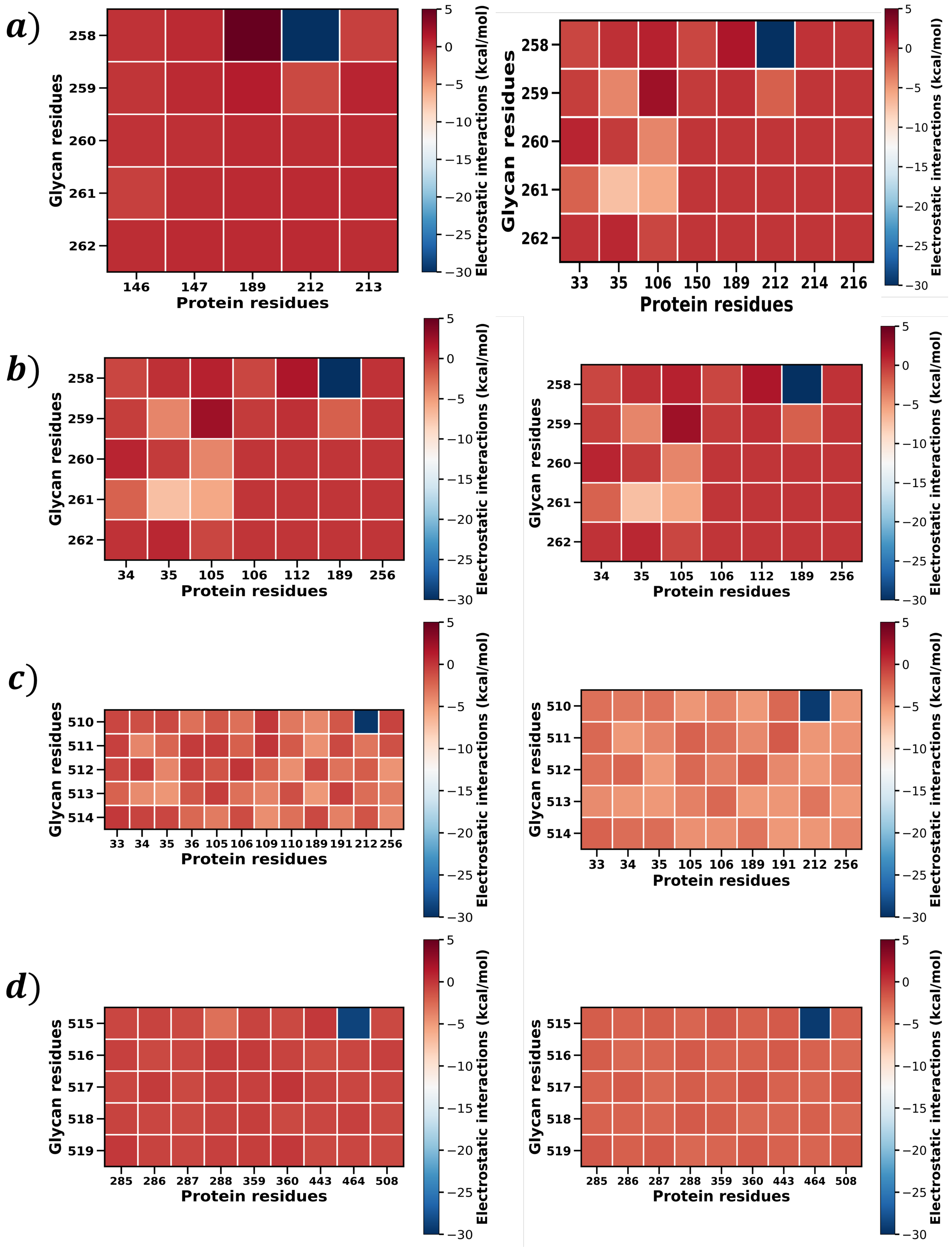

FIG. S5: Electrostatic component of linear interaction energy networks between the glycan chain and the surrounding protein residues corresponding to the highest free-energy barrier along the minimum free-energy pathway (MFEP). Panels (a), (b), and (c,d) depict the representative transition states connecting minima $7 \to 64$ in *G-mA*, $216 \to 1$ in *G-mB*, and $167 \to 8$ in *G-dAB*, respectively.

| S.No | Major Secondary Structure Elements | Constituent amino acid residues Chain A | | | | | Major Secondary Structure Elements | Constituent amino acid residues Chain B | | | | |
|---|---|---|---|---|---|---|---|---|---|---|---|---|
| | | Crystal Structure (PDB ID:3IAI) | *G-mA* | *U-mA* | *G-dA* | *U-dA* | | Crystal Structure (PDB ID:3IAI) | *G-mB* | *U-mB* | *G-dB* | *U-dB* |
| 01. | $\alpha$A | 23-26 | 23-26 | 23-25 | 23-26 | 23-26 | $\alpha$A | 23-26 | 23-26 | 23-25 | 23-25 | 23-26 |
| 02. | $\alpha$B | 28-31 | 28-31 | 28-31 | 28-30 | 28-31 | $\alpha$B | 28-31 | 32-34 | 28-31 | 28-31 | 28-31 |
| 03. | $\beta$a | 39-40 | 39-40 | 40 | 39-40 | 39-40 | $\beta$a | 39-40 | 39-40 | 38-40 | 39-40 | 39-40 |
| 04. | $\alpha$C | 42-44 | - | - | 42-44 | 42-44 | $\alpha$C | 42-44 | - | 42-44 | 42-44 | 42-44 |
| 05. | $\beta$b | 46-47 | 46-47 | 46-47 | 46-47 | 46-47 | $\beta$b | 46-47 | 46-47 | 46-47 | 46-47 | 46-47 |
| 06. | $\beta$**c** | 57 | 57-58 | 57-58 | - | 57-58 | $\beta$c | 54-54A | - | - | - | - |
| 07. | $\beta$A | 64-68 | 64-68 | 64-69 | 64-68 | 64-68 | $\beta$A | 64-68 | 64; 67-68 | 65-68 | 64-68 | 64-68 |
| 08. | $\beta$B | 76-79 | 72-79 | 72-79 | 76-79 | 76-79 | $\beta$B | 76-79 | 72-79 | 77-79 | 76-79 | 76-79 |
| 09. | $\beta$C | 86-90 | 87 | 86-88 | 85-90 | 86-90 | $\beta$C | 86-90 | 86-87 | 86-88 | 86-88 | 86-88 |
| 10. | $\beta$D | 93-102 | 95-102 | 95-102 | 93-102 | 93-102 | $\beta$D | 93-104 | 95-104 | 94-104 | 94-100 | 94-100 |
| 11. | $\beta$d | 115-116 | 115-116 | 115-116 | 115-116 | 115-116 | $\beta$d | 106 | - | 106 | - | - |
| 12. | $\beta$e | 119-120 | 119-120 | 119-120 | 119 | 119 | $\beta$e | 115-116 | 115-116 | 115-116 | 115-116 | 115-116 |
| 13. | $\beta$E | 123-132 | 123-132 | 123-132 | 123-132 | 123-132 | $\beta$f | 119-120 | 119-120 | 119-120 | 119 | 119 |
| 14. | $\alpha$D | 138-141 | 138-143 | 138-142 | - | 139-141 | $\beta$E | 123-132 | 123-132 | 123-132 | 123-132 | 123-132 |
| 15. | $\beta$F | 148-152 | 148-152 | 148-152 | 148-152 | 148-152 | $\alpha$D | 138-141 | 138-143 | 138-140 | 138-140 | 138-141 |
| 16. | $\alpha$E | 162-169 | 162-172 | 162-169 | 162-169 | 162-169 | $\beta$F | 148-156 | 148-157 | 148-156 | 148-151 | 148-151 |
| 17. | $\alpha$F | 171-173 | - | 171-174 | 171-173 | 171-174 | $\alpha$E | 165-169 | 165-172 | 165-173 | 164-169 | 162-173 |
| 18. | $\beta$G | 179-183 | 180-183 | 179-183 | 179-183 | 179-183 | $\alpha$F | 171-174 | - | - | 171-173 | - |
| 19. | $\beta$f | 187 | 186-187 | 186-187 | - | 186-187 | $\beta$G | 179-183 | 180; 183 | 179-182 | 179-183 | 180-183 |
| 20. | $\alpha$G | 188-190 | 188-189A | 188-189A | 189-190 | 188-189A | $\beta$g | 187 | 186 | 187 | 186 | - |
| 21. | $\beta$H | 198-201 | 198-201 | 198-201 | 198-201 | 198-201 | $\alpha$G | 188-190 | 188-190 | - | 188-190 | - |
| 22. | $\beta$I | 212-219 | 212-219 | 214-219 | 214-219 | 212; 214-219 | $\beta$H | 198-204 | 198-203 | 198-203 | 198-199 | 198-199 |
| 23. | $\beta$J | 223-225 | 223-224 | 223-225 | 223-225 | 223-225 | $\beta$I | 214-219 | 214-219 | 214-219 | 212-219 | 212-219 |
| 24. | $\alpha$H | 227-238 | 227-232 | 227-237 | 227-237 | 227-237 | $\beta$J | 223-225 | 223-226 | 223-225 | 223-226 | 223-225 |
| 25. | $\beta$g | 241 | - | 241 | 241 | 241-242 | $\alpha$H | 227-237 | 227-238 | 227-232 | 227-238 | 227-237 |
| 26. | $\beta$h | 247 | - | 247 | - | 246-247 | $\beta$h | 241-242 | - | 241 | 241-242 | 241-242 |
| 27. | - | - | - | - | - | - | $\beta$i | 246-247 | - | 247 | 246-27 | 246-247 |

TABLE S3: Major secondary structure elements and their constituent amino acid residues identified in the crystal and equilibrated structures of both monomeric and dimeric glycosylated and un-glycosylated HCA IX-c in aqueous solution, as determined using STRIDE.[1] The secondary structure assignments are based on the crystal structure of HCA IX (PDB ID: 3IAI).[2]

### REFERENCES

[1]M. Heinig and D. Frishman, Nucleic Acids Research **32**, 500 (2004).

[2]V. Alterio, M. Hilvo, A. Di Fiore, C. T. Supuran, P. Pan, S. Parkkila, A. Scaloni, J. Pastorek, S. Pastorekova, C. Pedone, *et al.*, Proceedings of the National Academy of Sciences USA **106**, 16233 (2009).

[3]R. Dey and S. Taraphder, The Journal of Physical Chemistry B **128**, 11054 (2024).

[4]G. Pérez-Hernández, F. Paul, T. Giorgino, G. De Fabritiis, and F. Noé, Journal of Chemical Physics **139**, 015102 (2013).

[5]M. M. Sultan and V. S. Pande, Journal of Chemical Theory and Computation **13**, 2440 (2017).

[6]S. V. Krivov and M. Karplus, The Journal of Chemical Physics **117**, 10894 (2002).

[7]D. Wales, *Energy Landscapes: Applications to Clusters, Biomolecules and Glasses*, Cambridge Molecular Science (Cambridge University Press, 2004).
